# The RNA demethylase FTO controls m^6^A marking on SARS-CoV-2 and classifies COVID-19 severity in patients

**DOI:** 10.1101/2022.06.27.497749

**Authors:** Lionel Malbec, Margot Celerier, Martin Bizet, Emilie Calonne, Heike Hofmann-Winkler, Bram Boeckx, Rana Abdelnabi, Pascale Putmans, Bouchra Hassabi, Lieve Naesens, Diether Lambrechts, Stefan Pöhlmann, Rachel Deplus, Leen Delang, Jana Jeschke, François Fuks

## Abstract

The RNA modification *N*^6^-methyladenosine (m^6^A) plays a key role in the life cycles of several RNA viruses. Whether this applies to SARS-CoV-2 and whether m^6^A affects the outcome of COVID-19 disease is still poorly explored. Here we report that the RNA demethylase FTO strongly affects both m^6^A marking of SARS-CoV-2 and COVID-19 severity. By m^6^A profiling of SARS-CoV-2, we confirmed in infected cultured cells and showed for the first time *in vivo* in hamsters that the regions encoding TRS_L and the nucleocapsid protein are multiply marked by m^6^A, preferentially within RRACH motifs that are specific to β-coronaviruses and well conserved across SARS-CoV-2 variants. In cells, downregulation of the m^6^A demethylase FTO, occurring upon SARS-CoV-2 infection, increased m^6^A marking of SARS-CoV-2 RNA and slightly promoted viral replication. In COVID-19 patients, a negative correlation was found between *FTO* expression and both SARS-CoV-2 expression and disease severity. FTO emerged as a classifier of disease severity and hence a potential stratifier of COVID-19 patients.

## Introduction

For over two years the world has been facing Severe Acute Respiratory Syndrome CoronaVirus 2 (SARS-CoV-2), the cause of Coronavirus Disease-2019 (COVID-19). The disease has spread at an alarming rate and has resulted (as of April 25, 2022) in more than 500 million confirmed cases and over 6 million deaths. About 20% of hospitalized COVID-19 patients develop a severe to critical pathology with acute respiratory failure. As they require prolonged oxygenation or ventilation (Wu & McGoogan, 2020; Stokes *et al*, 2020), intensive care units worldwide are overwhelmed. This situation has created an unprecedented international public health crisis that continues to threaten populations and the global economy, despite worldwide prevention measures and the progress of vaccine campaigns. To alleviate the effects of the pandemic, it is thus paramount to improve our understanding of COVID-19 severity and to identify markers for predicting and managing it. Currently, COVID-19 patients are stratified into mild, severe or critical cases on the basis on a set of parameters including CT images, oxygen saturation, and blood markers such as IL-6, CRP or lymphoid. (Zhou *et al*, 2020; Statsenko *et al*, 2021). This approach is laborious and biased, depends on the physician’s experience, and is applied when the patient is already symptomatic. We therefore need markers that allow a more efficient, unbiased stratification of COVID-19 patients, notably at earlier stages of the disease to help early intervention.

This is an area likely to be impacted by breakthroughs regarding the modifications that decorate RNA, highlighted as a crucial new regulatory layer(Frye *et al*, 2016). To date, over 150 distinct RNA modifications have been reported, *N*^6^-methyladenosine (m^6^A) being the most prevalent mark found in mRNA (Dominissini *et al*, 2012). Among eukaryotes, m^6^A is widely conserved and tends to occur within an RRACH consensus motif (where R represents G or A; H represents A, C, or U), especially in 3ʹ-untranslated regions (UTRs) near transcript stop codons. The mark is dynamically regulated, being installed by a multi-subunit complex composed of the METTL3-METTL14 heterodimer (Liu *et al*, 2014) and WTAP (Ping *et al*, 2014) and removed by the demethylases FTO and ALKBH5 (Jia *et al*, 2011; Zheng *et al*, 2013). Through YTH-family reader proteins (Wang *et al*, 2014, 2015; Shi *et al*, 2017), m^6^A regulates different aspects of mRNA metabolism (Roundtree *et al*, 2017; Song & Yi, 2020; Zhao *et al*, 2017; Yang *et al*, 2018) and impacts key pathophysiological processes (Chen & Wong, 2020; He *et al*, 2019). The mark and its machinery also play an essential role in the life cycles of several viruses of the families *Retroviridae* (HIV-1)(Kennedy *et al*, 2016; Lichinchi *et al*, 2016), *Polyomaviridae* (SV40)(Tsai *et al*, 2018), *Pneumoviridae* (HMPV)(Lu *et al*, 2020), *Flaviviridae* (IAV, HCV, ZIKV)(Courtney *et al*, 2017; Gokhale *et al*, 2016), *Herpesviridae* (KSHV)(Ye *et al*, 2017) and *Hepadnaviridae* (HBV)(Imam *et al*, 2018). Recent work has added *Coronaviridae* to this list, as SARS-CoV-2 transcripts have been found to undergo m^6^A-marking and as this appears to affect SARS-CoV-2 reproduction (Liu *et al*, 2021; Zhang *et al*, 2021a, 2021b; Li *et al*, 2021; Burgess *et al*, 2021; Campos *et al*, 2021). Upon SARS-CoV-2 infection, furthermore, changes in the expression and localization of the m^6^A machinery have been observed (Zhang *et al*, 2021b; Li *et al*, 2021). Yet these reports conflictingly attribute either an antiviral (replication-inhibiting) (Liu *et al*, 2021; Zhang *et al*, 2021a) or proviral (replication-favoring) (Zhang *et al*, 2021b; Li *et al*, 2021; Burgess *et al*, 2021) role to m^6^A, so the situation is far from clear. This warrants further investigation of the role of m^6^A in the SARS-CoV-2 viral cycle and in COVID-19 severity.

In the present study we have sought to better understand, in the light of clinical knowledge, how m^6^A and its machinery affect SARS-CoV-2 and its ability to cause severe COVID-19. We present data obtained on cultured cells, infected hamsters, and patient samples suggesting that (1) m^6^A deposited at highly conserved sites in SARS-Cov-2 and other pandemic coronaviruses may play an important role in the ongoing SARS-CoV-2 pandemic; (2) depletion of the m^6^A eraser FTO leads to elevated m^6^A marking of SARS-CoV-2 transcripts and to increased virus replication; and (3) FTO expression levels correlate inversely with both SARS-CoV-2 replication and COVID-19 severity. Collectively, our present work thus identifies m^6^A as a new regulator of SARS-CoV-2 infection and reveals FTO expression as a potential classifier of COVID-19 severity.

## Results

### The mark m^6^A decorates SARS-CoV-2 RNA both *in vitro* and *in vivo*

Recent reports suggest that SARS-CoV-2 RNA is marked (Kim *et al*, 2020) with m^6^A (Liu *et al*, 2021; Zhang *et al*, 2021a, 2021b; Li *et al*, 2021; Burgess *et al*, 2021; Campos *et al*, 2021), but inconsistencies regarding the positions and proposed role of m^6^A warrant further investigation. We thus performed m^6^A-specific antibody immunoprecipitation followed by high-throughput sequencing (m^6^A-seq) on total RNA extracted from SARS-CoV-2-infected Vero76 cells (**Fig. 1a**). Following read alignment with reference monkey and SARS-CoV-2 genomes and m^6^A peak calling, we first validated the reproducibility and reliability of our assay by testing the orthologous gene *SLC39A14* previously reported to be m^6^A-modified at its 3’UTR in mammals(Dominissini *et al*, 2012) (**Supplementary information, Fig. S1a**). We then identified three regions in the SARS-CoV-2 genome, grouped into two reproducible clusters, as corresponding to significant m^6^A peaks (**Fig. 1B** and **Supplementary information, Fig. S1b**). The largest m^6^A peak was located in the TRS_L region (**Fig. 1b**, **1c** and **Supplementary information, Fig. S1c**), responsible for programmed template switching by the viral RNA polymerase (RdRp) during viral sub-genomic RNA synthesis(Sola *et al*, 2015). Two additional m^6^A peaks were found in the region coding for the nucleocapsid protein (**Fig. 1b**, **1c** and **Supplementary information, Fig. S1c**), a key component facilitating viral mRNA transcription and replication, SARS-CoV-2 genome packaging into RNP viral particles, and RNA-interference-mediated antiviral responses(Masters, 2019; Cong *et al*, 2020; Mu *et al*, 2020). As regions identified by m^6^A-seq can contain one or multiple m^6^A residues, we next sought m^6^A motifs within our m^6^A-peak regions. We identified five m^6^A motifs in the TRS-L region and and eight in the nucleocapsid region, the consensus sequence RRACH appearing as the most prevalent motif (**Fig. 1d** and **Supplementary information, Fig. S1d**).

**Fig. 1.**
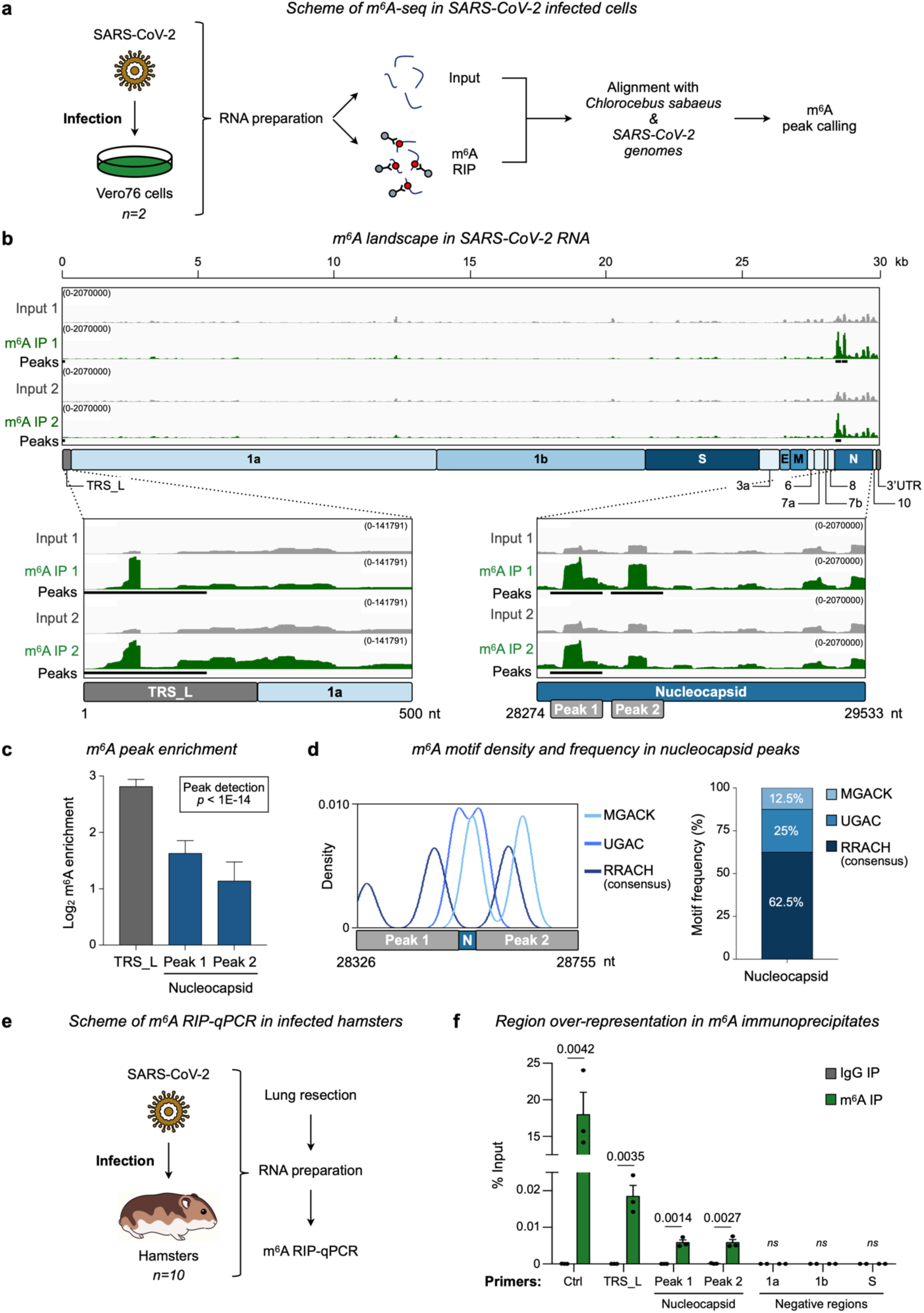
m^6^A decorates SARS-CoV-2 RNA *in vitro* and *in vivo*. **a** Scheme of m^6^A-seq in SARS-CoV-2-infected cells. Total RNA was extracted from SARS-CoV-2-infected Vero76 cells 24 h post-inoculation. Reads from Input and m^6^A RIP were aligned with STAR on both the *Chlorocebus sabaeus* and SARS-CoV-2 genomes. Peaks were then identified with m6aViewer. m^6^A-seq was performed in two independent experiments with technical duplicates. **b** m^6^A landscape in SARS-CoV-2 RNA. Integrative Genomics Viewer (IGV) tracks display reads from input (gray) and m^6^A IP (green) libraries along the SARS-CoV-2 RNA genome from two biological replicates. Three m^6^A peaks were identified from replicate 1 and two from replicate 2 (black squares). SARS-CoV-2 genome organization is indicated below. Enlarged views of the IGV tracks show the enrichment in reads corresponding to the SARS-CoV-2 TRS_L and nucleocapsid regions. **c** Bar plot shows m^6^A peaks in the TRS_L (gray) and nucleocapsid (blue) regions. The peaks detected exhibit a *p* value lower than 1E-14. **d** Density plot (left panel) displaying the distribution of known m^6^A motifs (RRACH, UGAC, and MGACK) across the m^6^A peaks of the nucleocapsid region. The stacked bar chart (right panel) shows motif frequencies within m^6^A peaks in the SARS-CoV-2 nucleocapsid region. **e** Scheme of m^6^A RIP-qPCR in infected hamsters. Ten hamsters were infected with SARS-CoV-2. Four days post-inoculation, the hamsters were sacrificed, their lungs collected, and total RNA extracted. RIP-qPCR was performed in three independent experiments to validate m^6^A peaks on SARS-CoV-2 RNA *in vivo*. **f** *In vivo* m^6^A peaks on SARS-CoV-2 viral RNA as determined by m^6^A RIP-qPCR. Lysates containing equal amounts of RNA extracted from infected hamster lungs were immunoprecipitated with either control IgG or m^6^A antibody. Primers amplifying the viral m^6^A-marked regions identified in monkey cells were used to assess immuno-precipitate enrichment in viral sequences. An m^6^A-marked oligo was used as positive control and regions found not to be marked in monkey cells were used as negative controls. Data represent means ± SEM. *P*-values were determined with a two-tailed unpaired Student’s t-test.

We then wondered whether m^6^A decorates SARS-CoV-2 RNA in the same regions *in vivo*, a question never addressed before in an animal model. We chose to use hamsters, previously shown to be highly permissive to SARS-CoV-2 infection and developing symptoms similar to those of COVID-19 patients(Kaptein *et al*, 2020) (**Fig. 1e**). When we performed m^6^A RIP-qPCR on total RNA extracted from lungs of infected hamsters, we observed the same three m^6^A peaks as *in vitro* (**Fig. 1f**). Our data thus indicate that SARS-CoV-2 RNA is marked with m^6^A in the TRS_L and nucleocapsid regions in both *in vitro* and *in vivo* models.

### The identified m^6^A-modified regions are conserved and specific to β-coronaviruses

SARS-CoV-2 belongs to the Coronaviridae family, which includes six additional coronaviruses known to have caused pandemics in humans (Cui *et al*, 2019): SARS-CoV (Drosten *et al*, 2003) and Middle East Respiratory Syndrome Coronavirus (MERS-CoV)(Zaki *et al*, 2012), which triggered severe pandemics preceding the present COVID-19 pandemic, and HCoV-HKU1(Woo *et al*, 2005), HCoV-OC43(Tyrrell & Bynoe, 1965), HCoV-NL63(van der Hoek *et al*, 2004), and HCoV-229E(Hamre & Procknow, 1966), which have become endemic. We wondered whether all these viruses might, like SARS-CoV-2, be marked by m^6^A in their TRS_L and nucleocapsid regions and whether this marking might play a role in their ability to cause a pandemic. To test this, we retrieved TRS_L and nucleocapsid sequences from all seven coronavirus strains and performed phylogenetic analyses. In line with a previous report(Chan *et al*, 2020), we found the TRS_L and nucleocapsid regions of SARS-CoV-2 to be phylogenetically closer to the corresponding regions of SARS-CoV (88%) than to those of the other coronaviruses (**Supplementary information, Fig. S2a** and **S2b**). On the basis of TRS_L and nucleocapsid sequence phylogeny, we further found the SARS-CoV-2, SARS-CoV, and MERS-CoV strains to cluster as a genus, appearing as a group within the β-coronavirus subtype. This highlights a potential evolutionary specificity of these regions (**Supplementary information, Fig. S2c** and **S2d**).

Next, we used SimPlot to assess the similarity of TRS_L and nucleocapsid sequences among human coronaviruses. We found the SARS-CoV-2 and SARS-CoV sequences to be almost 90% similar (**Fig. 2a** and **Supplementary information, Fig. S2e**). MERS-CoV and SARS-CoV-2 appeared nearly 50% identical to SARS-CoV-2 in the nucleocapsid region but not in the TRS_L region. In both regions, the other coronaviruses appeared less similar to SARS-CoV-2. Strikingly, our identified m^6^A peaks occurred in regions where the pandemic β-CoV strains appeared highly identical, suggesting that these sites have been evolutionarily conserved and that m^6^A might play an important role in the ability of these strains to become pandemic. This prompted us to investigate whether the m^6^A motifs identified within TRS_L and nucleocapsid sequences might be shared among human coronaviruses. Multiple sequence alignment revealed that several motifs within our three m^6^A peaks were indeed shared among human coronaviruses, but only among β-coronaviruses in general or pandemic β-coronaviruses in particular (**Fig. 2b** and **Supplementary information, Fig. S2f**).

**Fig. 2.**
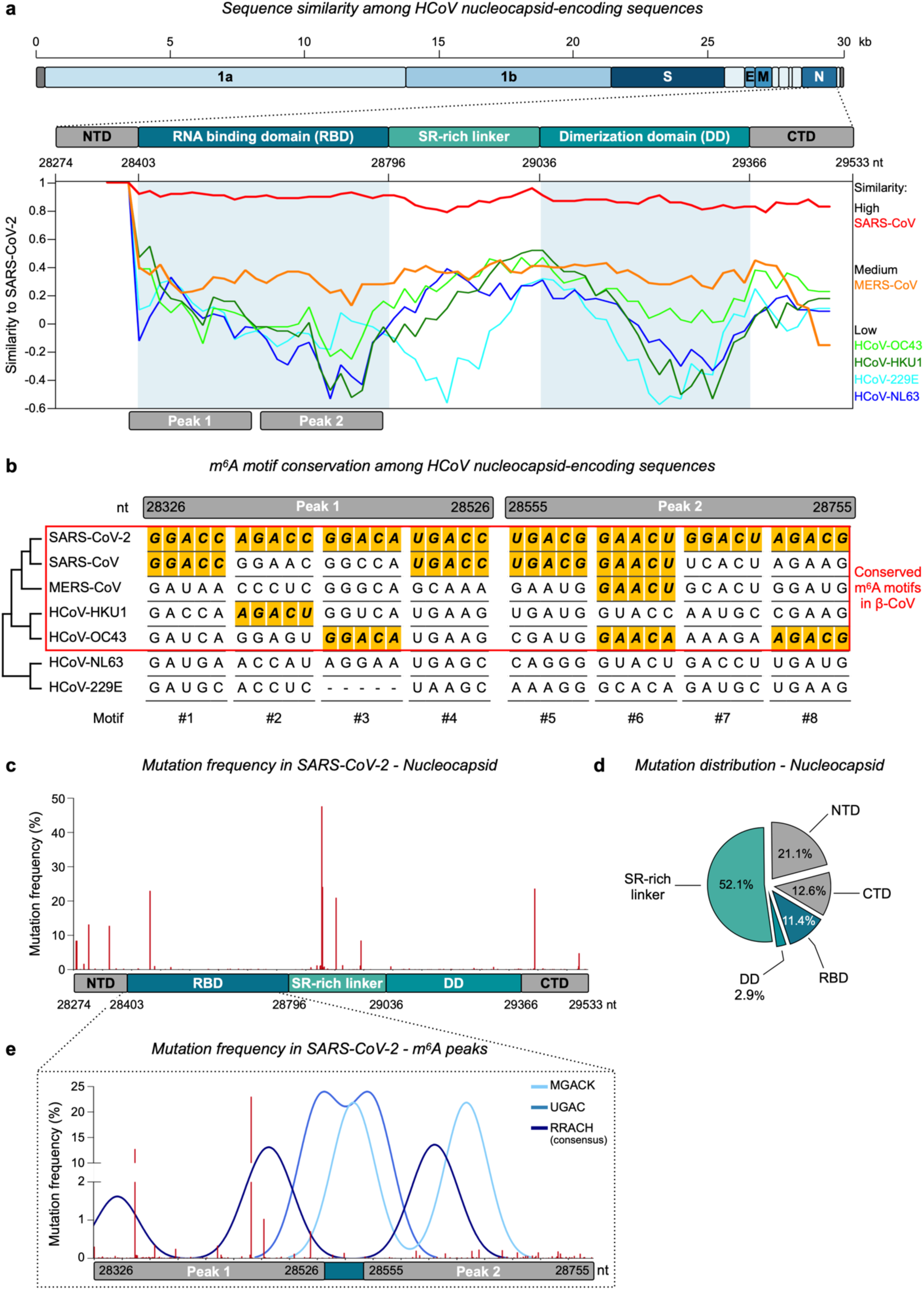
m^6^A sites are specific and conserved among SARS-CoV-2 strains. **a** Sequence similarity of nucleocapsid-protein-encoding regions of human coronaviruses. The upper diagram shows the genomic organization of the SARS-CoV-2 open reading frames (ORFs) and the domain organization of the nucleocapsid protein gene, based on NCBI sequence NC_045512.2. The SimPlot (lower panel) displays nucleotide similarities between SARS-CoV-2 and other human coronaviruses. It is based on the full-length sequence of the nucleocapsid gene derived from the multiple sequence alignment (MSA) of all human coronaviruses, with the following parameter settings: window 200 bp, step 20 bp, GapStrip: on, Kimura model. **b** MSA describing m^6^A consensus motif variations within peaks identified in SARS-CoV-2 across human coronavirus nucleocapsid sequences. The m^6^A consensus sequences conserved across HCoVs are colored in yellow. **c** Bar chart displaying mutation frequencies in nucleocapsid regions across SARS-CoV-2 variants as of April 25, 2022 from a total of 9,557,377 SARS-CoV-2 isolates with complete sequences. **d** Pie chart showing the distribution of mutations across the nucleocapsid domains of SARS-CoV-2 variants. **e** Bar chart (red) and density plot (blue) displaying, across SARS-CoV-2 variants, the frequencies and distribution of mutations in known m^6^A motifs (RRACH, UGAC and MGACK) associated with m^6^A peaks identified in the sequence encoding the nucleocapsid RNA binding domain.

Overall, our results support the view that conservation of m^6^A sites may have been important to phylogenetic evolution within the β-CoV genus. They suggest that m^6^A might play a role in the ongoing pandemic.

### The m^6^A sites are highly conserved among SARS-CoV-2 variants

We next focused on new and emerging SARS-CoV-2 variants to see if they might contain novel mutations in our identified m^6^A peaks. We downloaded a variation database grouping mutations in the TRS_L and nucleocapsid regions across SARS-CoV-2 variants from December 2019 to April 2022. About 88.7% of the isolates had been collected in developed countries (Europe and North America) (**Supplementary information, Fig. S2g**). Strikingly, among the 9.5 million isolates completely sequenced, the vast majority were variants of concern or interest, with Delta, Omicron and Alpha accounting, respectively, for 42.9%, 30.7% and 11.9% of the database (**Supplementary information, Fig. S2h**). Overall, the SARS-CoV-2 TRS_L and nucleocapsid regions showed very few high-frequency mutations (**Fig. 2c** and **Supplementary information, Fig. S2i**). In the nucleocapsid sequence, the few frequent mutations appeared within the central, intrinsically disordered serine/arginine-rich linker domain (SR), thought to be mutation-prone and crucial for the N protein’s RNA-mediated condensation during virion formation(Savastano *et al*, 2020). Other regions, including the RNA-binding domain (RBD) and dimerization domain (DD), showed very few mutations, suggesting a potential evolutionary pressure on these conserved folded structural domains essential to N protein function (**Fig. 2d**).

We next focused specifically on variations within the regions corresponding to identified m^6^A peaks and motifs. Overall, we found the m^6^A**-**peak regions to be highly conserved in both the TRS_L and nucleocapsid sequences (**Fig. 2e** and **Supplementary information, Fig. S2j**). The RRACH, UGAC, and MGACK motifs appeared conserved as well, although we did observe one significant high-frequency event overlapping with one of the potentially methylated RRACH motifs in the nucleocapsid RBD region.

Our m^6^A peaks identified in SARS-CoV-2 are thus located within highly conserved regions that are rarely affected by new variations. This strengthens the view that m^6^A may play an important role in the ongoing COVID-19 pandemic.

### *FTO* silencing is linked to increased m^6^A marking of SARS-CoV-2 RNA and to impaired virus replication

We next investigated whether SARS-CoV-2 might affect its own m^6^A marking by regulating the host’s m^6^A machinery. In Vero76 cells, we measured levels of m^6^A methylase (METTL3, METTL14, WTAP) and m^6^A demethylase (FTO, AKLBH5) mRNAs before and after infection. Twenty-four hours post-infection, we observed a slight but significant decrease in METTL3, METTL14, and WTAP mRNAs and a more pronounced reduction in FTO but not ALKBH5 mRNAs (**Fig. 3a** and **Supplementary information, Fig. S3a**). These findings seem to be in agreement with a previous report revealing decreased FTO protein levels 24 h and 48 h post-infection(Zhang *et al*, 2021b), pointing to FTO as a potential key regulator of m^6^A in SARS-CoV-2. In line with the weak reduction we observed in transcript-level *METTL3* and *METTL14* expression, this same report found the corresponding protein levels to be unchanged 24h post-infection (Zhang *et al*, 2021b).

**Fig. 3.**
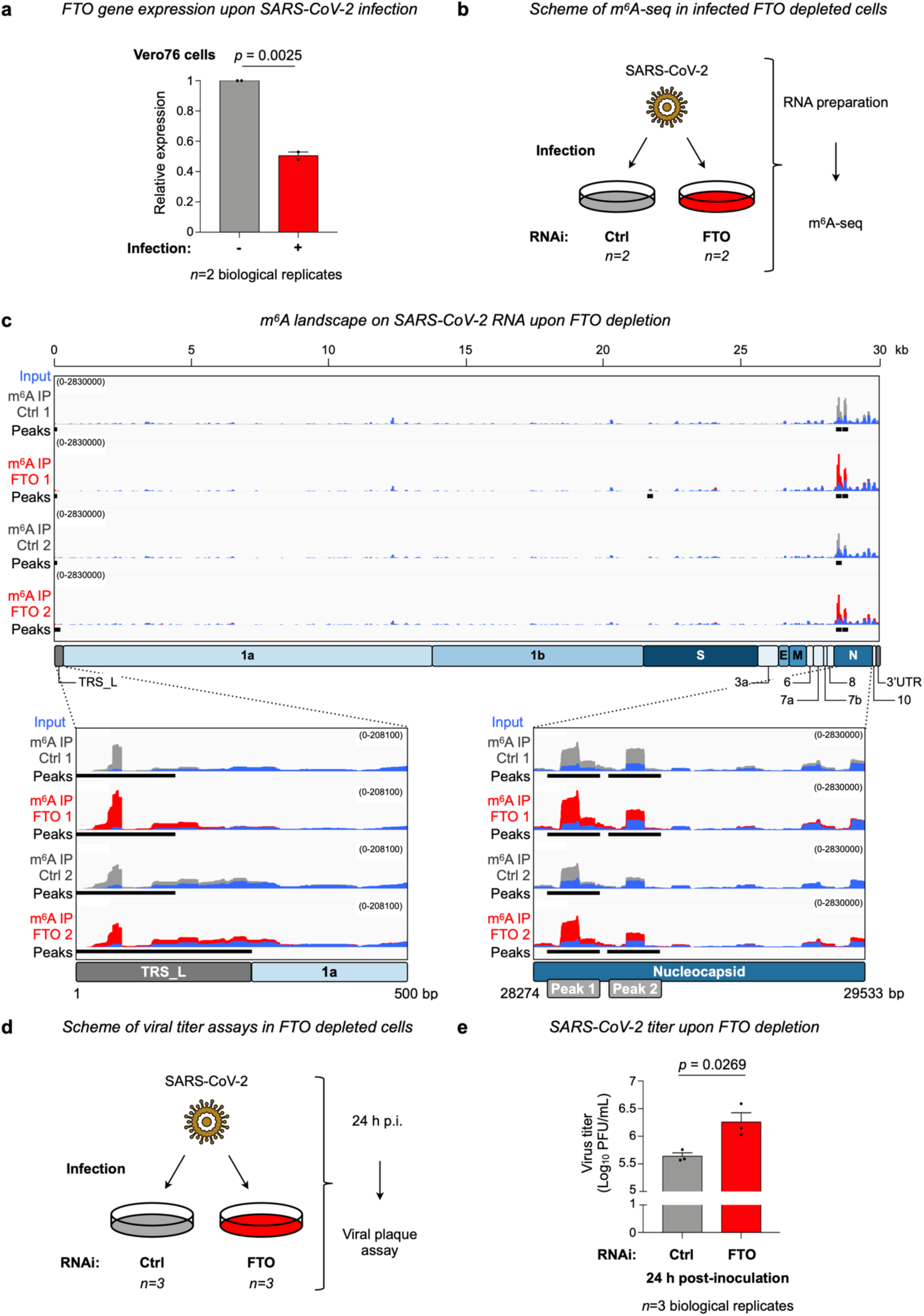
The m^6^A demethylase FTO impairs SARS-CoV-2 replication. **a** *FTO* gene expression upon SARS-CoV-2 infection in monkey cells. RT-qPCR assays show reduced levels of FTO mRNA in Vero76 cells after SARS-CoV-2 infection. Dots represent mean expression in independent experiments conducted with technical triplicates. **b** Scheme of m^6^A-seq in infected FTO-depleted monkey cells. Total RNA was extracted from SARS-CoV-2-infected control (Ctrl) and FTO-depleted Vero76 cells 24 h post-inoculation. Then m^6^A-seq was performed in two independent experiments with technical duplicates. **c** m^6^A landscape of SARS-CoV-2 RNA upon FTO depletion. IGV tracks display input reads (blue) and m^6^A IP reads from control (gray) and FTO-depleted (red) libraries along the SARS-CoV-2 RNA genome. Three m^6^A peaks were identified on the genome of Ctrl cells and five on that of FTO-depleted cells (black squares). The SARS-CoV-2 genome organization is indicated below. Enlarged views of the IGV tracks show the m^6^A peaks within the SARS-CoV-2 TRS_L and nucleocapsid regions. **d** Scheme of viral titer assays on infected FTO-depleted monkey cells. SARS-CoV-2 replication was assessed by viral plaque assay on infected control and FTO-depleted Vero76 cells and performed in three independent experiments with technical triplicates. **e** Bar chart showing increased SARS-CoV-2 replication in Vero76 cells upon FTO depletion 24 h post-inoculation. Plaque-forming units (PFU) of infectious SARS-CoV-2 particles in culture supernatants were determined by plaque titration with Vero-TMPRSS2 cells. Dots represent mean titers for three independent experiments conducted with technical triplicates. Data represent means ± SEM. *P*-values were determined with a two-tailed unpaired Student’s t-test.

To study whether FTO regulates m^6^A in SARS-CoV-2, we silenced the *FTO* gene in Vero76 cells, using short hairpin RNA interference (RNAi). We first confirmed successful depletion of FTO at the RNA and protein levels (**Supplementary information, Fig. S3b** and **S3c**). We then performed m^6^A-seq, using RNA collected from SARS-CoV-2-infected RNAi Ctrl and RNAi FTO cells (**Fig. 3b**). In line with our initial findings (cf. Fig. 1b), we detected m^6^A peaks in the TRS_L and nucleocapsid regions of SARS-CoV-2 RNA (**Fig. 3c**). In FTO-depleted cells, furthermore, we discovered two additional peaks, one in the TRS_L region and one in the spike region (**Supplementary information, Fig. S3d**). Importantly, m^6^A occupancy appeared increased in the TRS_L and nucleocapsid regions in FTO-depleted versus control cells (**Fig. 3c**). This strongly suggests that FTO regulates m^6^A in SARS-CoV-2.

We next assessed the impact of m^6^A on the SARS-CoV-2 viral cycle. For this we performed viral plaque assays on RNAi Ctrl and RNAi FTO Vero76 cells inoculated with equal amounts of SARS-CoV-2 (**Fig. 3d** and **Supplementary information, Fig. S3e**). We observed, 24 h post-inoculation, a slight but significant increase in virus titer in *FTO*-silenced versus control cells (**Fig. 3e**). This suggests that FTO, via m^6^A, plays a role in SARS-CoV-2 replication.

Taken together, our findings suggest that SARS-CoV-2 infection *in vitro* correlates with downregulation of the m^6^A eraser FTO, increased m^6^A marking of the SARS-CoV-2 RNA in the TRS_L and nucleocapsid regions, and increased SARS-CoV-2 reproduction.

### In patients, FTO downregulation correlates inversely with SARS-CoV-2 infection and COVID-19 severity

Next, we investigated the clinical relevance of our findings. As the above data indicate that FTO, via m^6^A, may impact the SARS-CoV-2 viral cycle, we wondered whether this might affect the course of COVID-19 in patients. To test this, we retrieved single-cell RNA-seq data from bronchioalveolar lavage (BAL) samples collected from 44 hospitalized patients with pneumonia. Of these patients, 31 tested positive for SARS-CoV-2 by RT-qPCR applied to a nasopharyngeal swab or to a lower respiratory tract sample. They are referred to as the ‘COVID-19’ group. Thirteen of them tested negative and are referred to as the ‘control’ group(Wauters *et al*, 2021) (**Fig. 4a**). The available data concerned 31,923 cells from COVID-19 control samples and 84,874 cells from COVID-19 samples. COVID-19 patients were further stratified on the basis of disease severity (oxygenation needs) into 5 ‘mild’ (n= 9,998 cells) and 26 ‘severe’ (n= 74,876 cells) cases. Cluster analysis based on UMAP assigned the 116,797 cells to different cell-type clusters, including lymphoid, myeloid, and epithelial cells (**Supplementary information, Fig. S4a**), confirming a recent study (Wauters *et al*, 2021). Myeloid cells were the most abundant cell type, amounting to 55.2% (n=64,485) of total cells. Lymphoid and epithelial cells amounted, respectively, to 25.8% (n=30,097) and or 19% (n=22,215) of total cells. Interestingly, FTO expression appeared highest in epithelial cells. The proportion of lymphoid cells was higher, and those of myeloid and epithelial cells were lower, in COVID-19 vs control samples (**Supplementary information, Fig. S4b**). We then sub-clustered epithelial cells into secretory, basal, ciliated, squamous, inflammatory, and AT2 epithelial cells. We found FTO to be expressed mainly in the most abundant epithelial cell types, especially ciliated cells (**Supplementary information, Fig. S4c**). We observed changes in epithelial cell lineage composition according to the disease status (**Supplementary information, Fig. S4d**): samples from COVID-19 patients were poorer than control samples in AT2, secretory, and squamous epithelial cells and richer in ciliated and basal cells. Together, these analyses revealed that FTO is predominantly expressed in ciliated lung epithelial cells, identified as the main target for SARS-CoV-2 entry(Ziegler *et al*, 2021) and whose relative abundance appears altered in COVID-19 patients.

**Fig. 4.**
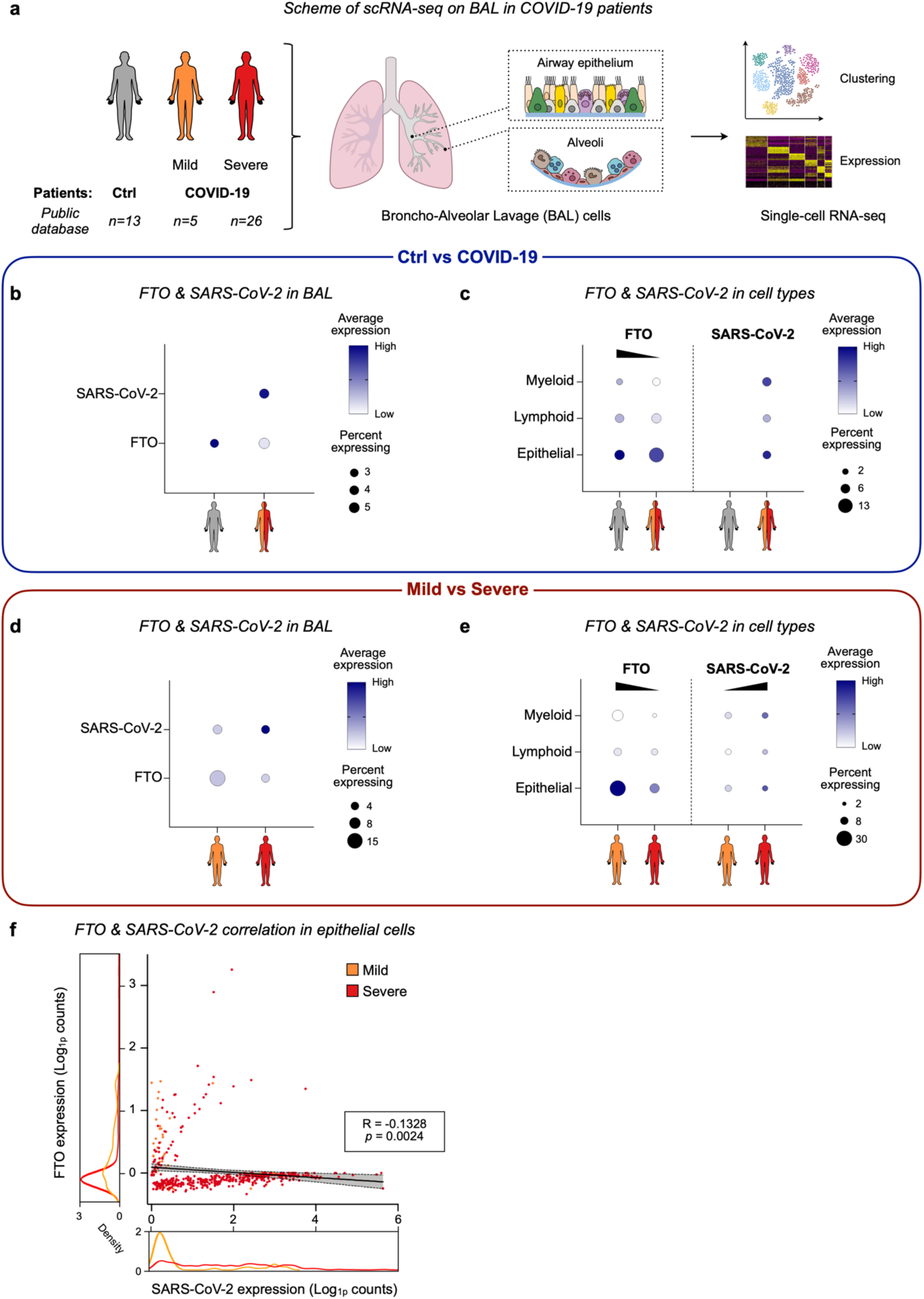
*FTO* downregulation correlates with higher SARS-CoV-2 expression. **a** Scheme of single-cell RNA-seq (scRNA-seq) in COVID-19 patients (Wauters *et al*, 2021). The 44 patients of this cohort included 13 controls (Ctrl) and 31 COVID-19 patients (5 with mild and 26 with severe symptoms). Lung cells from the airway epithelium and alveoli were collected by broncho-alveolar lavage (BAL) and subjected to scRNA-seq analysis **b, c** Dot plots show the differential expression of *FTO* and SARS-CoV-2 between Ctrl and COVID-19 patient cells in the entire BAL dataset (b) and in the myeloid, lymphoid and epithelial cell populations identified in BAL samples (c). **d, e** Dot plots display the differential expression of the *FTO* gene and of SARS-CoV-2 between cells from patients with mild and severe COVID-19 in the entire BAL dataset (d) and in the identified myeloid, lymphoid, and epithelial cell subsets (e). In the dot plots, dot size represents the proportion of cells expressing the gene of interest within the cell type considered and dot color represents the average gene expression level within that cell type. **f** *FTO* and SARS-CoV-2 expression are anti-correlated in epithelial cells from COVID-19 patients. Scatter plot displaying *FTO* and SARS-CoV-2 expression for each epithelial cell from COVID-19 patients with mild or severe disease. Density plots illustrate the distribution of gene expression among cell types. The one-tailed Pearson’s test was used for the correlation analyses.

Then, we investigated the relationship between *FTO* expression and SARS-CoV-2 in the BAL data set. As expected, and taking all cells into account, we detected SARS-CoV-2 in the COVID-19 but not the control samples (**Fig. 4b**). *FTO* expression appeared strongly decreased in COVID-19 as compared to control samples, although the fraction of cells expressing *FTO* remained unchanged. This corroborates the inverse correlation revealed between *FTO* expression and SARS-CoV-2 replication *in vitro* (cf. Fig. 3e). Observations on the distinct BAL cell types were similar (**Fig. 4c**): SARS-CoV-2 was detected exclusively in myeloid, lymphoid, and epithelial cells from COVID-19 samples, while these cells displayed lower *FTO* expression in COVID-19 than in control samples. It is worth noting that the COVID-19 samples displayed a slightly larger fraction of *FTO*-expressing epithelial cells than the control samples, but a lower overall level of *FTO* expression.

Lastly, we studied *FTO* and SARS-CoV-2 ORF expression with respect to COVID-19 severity. In BAL samples, we found *FTO* expression to be slightly lower in severe vs mild cases of COVID-19. The respective proportions of *FTO*-expressing cells were 4% and 16%. SARS-CoV-2 expression was increased in severe samples, although the fraction of infected cells was the same in mild and severe cases (**Fig. 4d**). Similar results were obtained for the distinct BAL cell types (**Fig. 4e**). Strikingly, the greatest difference in *FTO* expression between severe and mild cases was observed in epithelial BAL cells, found to express *FTO* most strongly. To further explore this finding, we focused on the lung epithelial cell lineages found to express *FTO* most abundantly (**Supplementary information, Fig. S4c**) and reported to be most permissive to SARS-CoV-2 infection(Ziegler *et al*, 2021). We found lower *FTO* expression in AT2, ciliated, squamous, and secretory cells from COVID-19 vs control samples and from severe vs mild cases (**Supplementary information, Fig. S4e** and **S4f**). On the other hand, these same cell types showed SARS-CoV-2 ORF expression only in COVID-19 samples, and this expression was higher in samples from severe versus mild cases. These results suggest an inverse relationship between FTO expression and SARS-CoV-2 infection that seems to be linked to COVID-19 severity in patients. We thus measured the correlation between FTO and SARS-CoV-2 expression in mild and severe cases. This revealed a significant anti-correlation depending on severity status (**Fig. 4f**): mild and severe COVID-19 cases exhibited distinct expression profiles, mild cases having higher FTO and lower SARS-CoV-2 expression and severe cases having lower FTO and higher SARS-CoV-2 expression.

These results for patient samples thus reveal an inverse correlation between FTO expression and SARS-CoV-2 infection or COVID-19 severity. They point, together with our *in vitro* data, to a potential role for FTO-mediated m^6^A demethylation in determining the course of COVID-19.

### FTO expression classifies COVID-19 severity in patients

We next examined the potential clinical utility of the observed anti-correlation between FTO expression and COVID-19 severity. Using a comprehensive machine learning approach (see Methods), we first assessed in epithelial cells of the BAL data set whether FTO expression might predict COVID-19 severity in COVID-19 patients (Wauters *et al*, 2021) (**Fig. 5a**). “Receiver Characteristic Operator” (ROC) curves revealed an “area under the curve” (AUC) of 0.768 (95% confidence interval (CI), 0.704 - 0.833), suggesting that FTO expression can distinguish mild from severe cases with a mean accuracy of 76.8%. To benchmark the prediction accuracy of FTO expression, we compared it with several reported biomarkers of COVID-19 severity (Mick *et al*, 2020; Merad & Martin, 2020). For IFI6, IL1B, IL1R2, CCR2, CCR5, and IL6 we observed respective AUCs of 0.732, 0.509, 0.476, 0.519, 0.523, and 0.493, indicating a lower accuracy than measured for FTO (**Fig. 5b** and **Supplementary information, Fig. S5a**). We next combined the two markers with the highest accuracy, FTO and IFI6 expression, and found a slightly improved predictive accuracy with this 2-gene signature (AUC 0.793, 95% CI between 0.718 to 0.869). This suggests that the information conveyed by the two genes is complementary, albeit partially overlapping (**Fig. 5c**). Together, these findings reveal FTO expression as a reliable classifier of COVID-19 severity, with 70.4–83.3% accuracy, when applied to BAL samples from hospitalized COVID-19 patients.

**Fig. 5.**
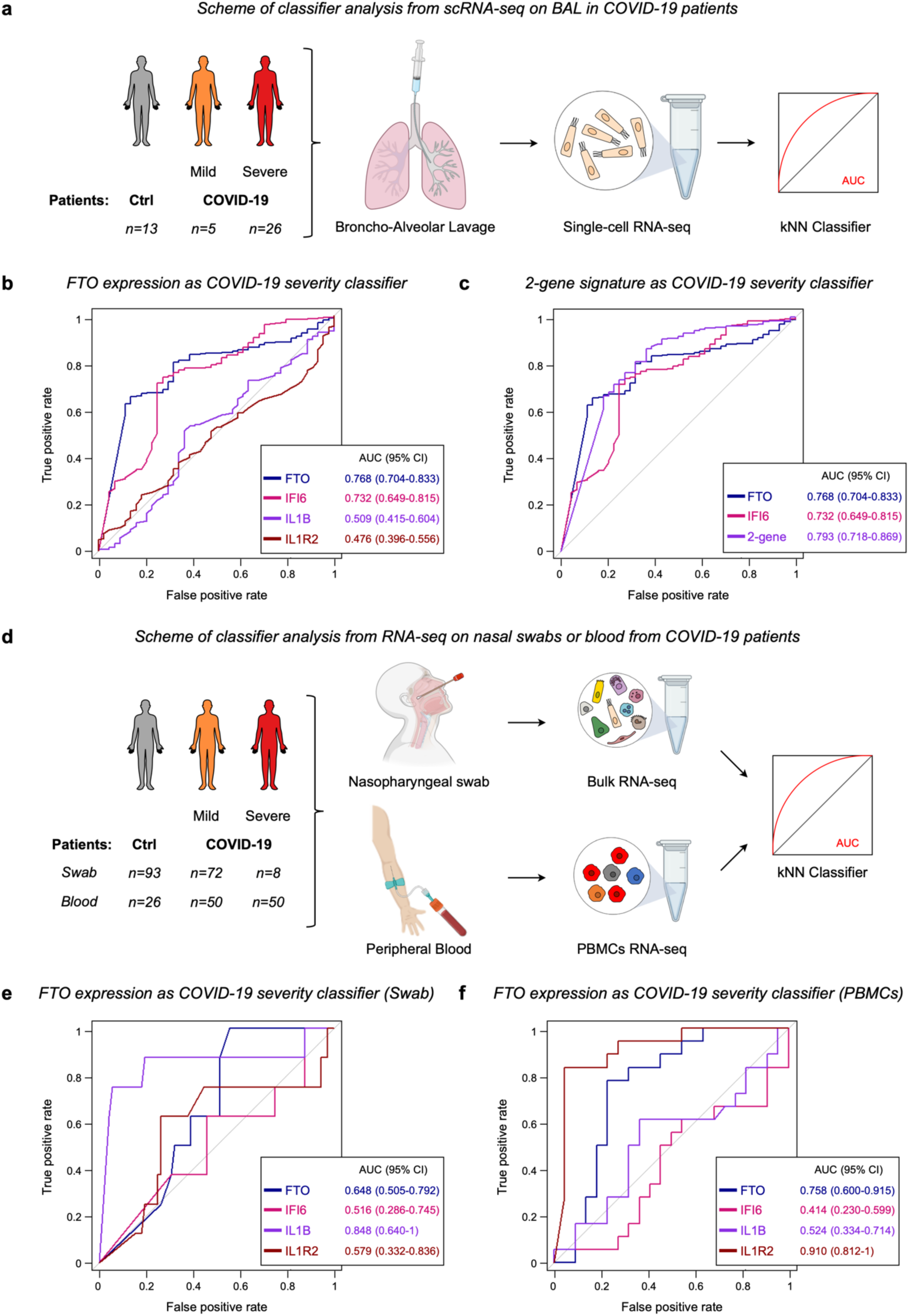
*FTO* expression in patient samples classifies COVID-19 severity. **a** For classifier construction, the approach used was k-Nearest Neighbor (kNN) machine learning on a single-cell RNA-seq dataset of gene expression levels in lung epithelial cells obtained from a patient cohort of 13 control (Ctrl) and 31 COVID-19 patients (5 with mild and 26 with severe disease) subjected to broncho-alveolar lavage (BAL)(Wauters *et al*, 2021). **b** Receiver operating characteristic (ROC) curves show the performance of *FTO* expression as a diagnostic classifier of COVID-19 severity as compared to *IFI6*, *IL1B*, and *IL1R2* gene expression in patient epithelial cells from BAL samples. **c** ROC curves showing the ability of a 2-gene signature (FTO plus IFI6) in BAL samples to classify COVID-19 severity. **d** For classifier construction, a kNN machine learning approach was used on an RNA-seq dataset obtained from a cohort of 93 control (Ctrl) and 80 COVID-19 patients (72 with mild and 8 with severe disease) subjected to nasal swab sampling(Ng *et al*, 2021) and from a cohort of 26 control (Ctrl) and 100 COVID-19 patients (50 with mild and 50 with severe disease) subjected to blood sampling(Overmyer *et al*, 2021). **e**, **f** ROC curves show the performance of *FTO* expression in patient cells from nasal swab samples (e) and PBMCs (f) as a diagnostic COVID-19 severity classifier, as compared to *IFI6*, *IL1B*, and *IL1R2* gene expression (e) and PBMCs (f). The mean of the area under the ROC curve (AUC) and the 95% confidence interval are indicated.

To assess COVID-19 severity and optimize patient triage and resource allocation upon admission to hospital, physicians routinely use nasal swab or blood testing (World Health Organization, 2021). Therefore, our next question was whether *FTO* expression, determined by RNA-seq applied to nasopharyngeal swabs(Ng *et al*, 2021) or blood samples(Overmyer *et al*, 2021) from COVID-19 patients might serve as classifier of COVID-19 severity (**Fig. 5d**). We found it to serve this purpose with a mean accuracy of 64.8% (50% - 79.2%) for nasopharyngeal swabs and 75.8% (CI, 60% - 91.5%) for blood samples (**Fig. 5e**, **5f** and **Supplementary information, Fig. S5b**, **S5c**). With nasal swabs, levels of some reported markers of COVID-19 severity (IFI6, IL1R2, CCR5 and IL6) proved unable to distinguish mildly from severely affected patients, whereas IL1B and CCR2, did so effectively, with respectively 84.8% (CI, 64% - 100%) and 90% (CI, 81.7% - 98.3%) accuracy (**Fig. 5e and Supplementary information, Fig. S5b**). In blood, only levels of IL1R2 and CCR5 could distinguish patients according to COVID-19 severity, with AUCs of 91% (CI, 81.2% - 100%) and 85.1% (CI, 73.5% - 95.7%), respectively (**Fig. 5f and Supplementary information, Fig. S5c**). Thus, FTO is the only marker found to predict COVID-19 severity reliably in both swab and blood samples of COVID-19 patients.

Together, our results identify *FTO* expression as a new and marker for determining COVID-19 severity, that can be used with different types of samples.

## Discussion

Internal marking with m^6^A has been shown to affect both the replication and spread of cytoplasmic RNA viruses such as HIV, IAV, and HBV(Kennedy *et al*, 2016; Lichinchi *et al*, 2016; Tsai *et al*, 2018; Courtney *et al*, 2017; Gokhale *et al*, 2016; Imam *et al*, 2018; Ye *et al*, 2017; Lu *et al*, 2020). Yet our understanding of whether and how m^6^A impacts the SARS-CoV-2 life cycle and COVID-19 pathogenesis is still limited. Recent reports suggest that SARS-CoV-2 hijacks the host m^6^A machinery to modify its own RNA (Liu *et al*, 2021; Zhang *et al*, 2021a, 2021b; Li *et al*, 2021; Burgess *et al*, 2021; Campos *et al*, 2021), but the consequence of this remains controversial, some studies suggesting that m^6^A decreases (Liu *et al*, 2021; Zhang *et al*, 2021a) and others that it increases SARS-CoV-2 abundance (Zhang *et al*, 2021b; Li *et al*, 2021; Burgess *et al*, 2021). How the host m^6^A machinery affects COVID-19 severity is also unclear (Li *et al*, 2021; Meng *et al*, 2021). In the present study we validate and extend previous findings showing the existence of m^6^A in the SARS-CoV-2 RNA genome. We identify FTO as an m^6^A demethylase of SARS-CoV-2 RNA and as a previously unrecognized regulator and marker of COVID-19 severity.

Recent studies using different m^6^A profiling approaches on human and monkey cells have revealed m^6^A sites in SARS-CoV-2 RNA, especially at the 3ʹ end of the viral genome (Liu *et al*, 2021; Zhang *et al*, 2021a, 2021b; Li *et al*, 2021; Burgess *et al*, 2021; Campos *et al*, 2021). In support of these findings, we have observed m^6^A peaks in the TRS_L and nucleocapsid (N) regions of SARS-CoV-2 isolated from monkey cells, further identifying the RRACH consensus sequence as the most prevalent motif within these peaks. We have also found the existence of these m^6^A sites on SARS-CoV-2 in a hamster model. The m^6^A sites identified here and in recent studies in the viral TRS_L and N regions thus appear as high-confidence m^6^A sites that could be key targets for characterizing the role of viral m^6^A in the life cycle of SARS-CoV-2. In this regard the nucleocapsid region is of particular interest: the fact that it was highlighted in all studies suggests that m^6^A may have a crucial regulatory function in this region. Our study, however, has not confirmed all of the m^6^A sites observed in other studies. A combination of factors might explain such discrepancies: different resolutions achieved by different m^6^A mapping technologies, different experimental conditions (e.g. timing after SARS-CoV-2 infection), different cell lines and SARS-CoV-2 strains used. Thus, the existence and exact location of most SARS-CoV-2 m^6^A sites remain uncertain. This warrants further investigation, beyond the efforts already made in this direction (Liu *et al*, 2021; Burgess *et al*, 2021; Campos *et al*, 2021).

We have further studied the evolutionary conservation of the SARS-CoV-2 genomic regions identified here as marked by m^6^A (TRS_L and N). We find them to be relatively conserved and specific to the genus β-coronavirus. Interestingly, the β-coronaviruses having caused recent pandemics, i.e. SARS-CoV-2 and SARS-CoV especially but also MERS-CoV, show a similar number of conserved m^6^A sites with particularly high m^6^A motif sequence similarity. These findings point to evolutionary conservation of m^6^A sites within the β-CoV genus, likely to reflect an important role for m^6^A in pandemic spread. Accordingly, we have found the identified m^6^A sites to remain mostly unaffected in SARS-CoV-2 variants. Yet there is at least one position, within the N region, where we have found a mutation liable to negatively affect m^6^A methylation by disrupting the RRACH motif. Whether differences in m^6^A methylation between SARS-CoV-2 variants are linked to differences in transmission or pathogenicity thus deserves further study.

Our findings provide crucial support to the idea that m^6^A is dynamically regulated in the SARS-CoV-2 RNA genome. On the one hand, we show that when the FTO gene is silenced, the m^6^A level increases in SARS-CoV-2 RNA. We further show that FTO depletion promotes SARS-CoV-2 replication in monkey cells. This suggests that FTO may participate in regulating the SARS-CoV-2 life cycle and that m^6^A may have a pro-viral function in SARS-CoV-2. Accordingly, we reveal an inverse correlation between FTO expression and SARS-CoV-2 infection in patient samples. Although m^6^A has previously been implicated in the regulation of SARS-CoV-2, whether it promotes or inhibits SARS-CoV-2 replication has remained unclear (Liu *et al*, 2021; Zhang *et al*, 2021a, 2021b; Li *et al*, 2021; Burgess *et al*, 2021). In agreement with our study, the majority of reports favor a pro-viral role of m^6^A in SARS-CoV-2 (Zhang *et al*, 2021b; Li *et al*, 2021; Burgess *et al*, 2021). This is, notably, also in agreement with findings on other RNA viruses such as IAV, HIV, and ZIKV (Kennedy *et al*, 2016; Courtney *et al*, 2017; Gokhale *et al*, 2016). Why do some reports conclude that m^6^A plays an anti-viral role by hindering the SARS-CoV-2 viral cycle (Liu *et al*, 2021; Zhang *et al*, 2021a)? It could be that m^6^A affects SARS-CoV-2 RNA in different ways via different mechanisms, as observed for mammalian RNA (Roundtree *et al*, 2017). Alternatively, discrepant findings might reflect differences in m^6^A methylation patterns due to different cell line models, different SARS-CoV-2 strains, and/or different timing post-infection. These potential sources of divergence should be taken into account in future studies aiming to clarify the role of m^6^A in SARS-CoV-2 infection. This is critical to propose strategies targeting m^6^A and its machinery in the fight against COVID-19 (Liu *et al*, 2021; Meng *et al*, 2021). Future studies should focus on further identifying and characterizing functional m^6^A sites in SARS-CoV-2 RNA to ascertain whether it’s the viral m^6^A or the host m^6^A that regulates SARS-CoV-2 abundance. We and others have found SARS-CoV-2 infection to affect the expression and localization of the host m^6^A machinery, causing changes in both the viral and host m^6^A methylomes (Liu *et al*, 2021; Zhang *et al*, 2021b; Meng *et al*, 2021). Linking changes in the host m^6^A methylome to altered immune responses, Meng et al. and Li et al. suggest a crucial role for host m^6^A in regulating SARS-CoV-2 abundance (Li *et al*, 2021; Meng *et al*, 2021). It is worth noting that we and others (Liu *et al*, 2021; Zhang *et al*, 2021a, 2021b; Li *et al*, 2021) have worked on Vero cells, which are immunodeficient. The data obtained on these cells thus support the view that viral m^6^A is important in regulating SARS-CoV-2 abundance. While more in-depth studies are needed to better understand this regulation and decipher its mechanisms, our results already suggest that targeting m^6^A could be an effective new treatment strategy for SARS-CoV-2 infection, affecting both the innate immune response and the viral life cycle.

The most striking findings of our study concern the m^6^A demethylase FTO. We show that (1) FTO expression is reduced upon SARS-CoV-2 infection; and (2) there is an inverse correlation between the FTO expression level and disease severity in patients. These findings tally with those of recent studies showing increased m^6^A abundance in the RNA of SARS-CoV-2-infected cells (Zhang *et al*, 2021b; Li *et al*, 2021) and a positive correlation of this increase with COVID-19 severity in patients (Meng *et al*, 2021). Together, these data point to the m^6^A demethylase FTO as a potential regulator and predictor of COVID-19 severity. With our BAL data set, we find FTO expression to distinguish mild from severe COVID-19 cases, with an accuracy higher than those of existing markers used to stratify COVID-19 severity (Gallo Marin *et al*, 2021). Our tests on nose swabs and blood samples further validate FTO expression as a marker of COVID-19 severity. Although FTO does not reach, in these samples, the accuracy of some existing inflammatory markers, it appears as the only marker that can reliably predict COVID-19 severity across the different sample types. Thus, our study identifies FTO expression as a new and universal marker of COVID-19 severity. It came as a surprise to us that this marker has predictive value in blood samples. This suggests that FTO may regulate SARS-CoV-2 spread and COVID-19 severity not only via epithelial cells. Accordingly, the findings of a very recent study suggest that monocytes and macrophages, in addition to lung epithelial cells, can undergo SARS-CoV-2 infection (Junqueira *et al*, 2022). Here we show that FTO expression stratifies COVID-19 patients for disease severity when they are already symptomatic and admitted to hospital. Whether it might predict severe outcome at an earlier stage of the disease, thereby allowing early intervention, remains to be tested in appropriate cohorts (not available for this study).

In summary, our work substantiates the existence and dynamic regulation of m^6^A in SARS-CoV-2 RNA and reveals the m^6^A demethylase FTO as a potential new regulator and marker of COVID-19 severity. Our findings both enhance our understanding of COVID-19 pathology and offer encouraging prospects for the development of novel patient triage approaches based on m^6^A and its machinery.

## Materials and Methods

### SARS-CoV-2 strains

All virus-related experiments were conducted in BSL3 core facilities, at either the German Primate Center (Germany) or the Rega Institute (Belgium). The SARS-CoV-2 strain used to infect Vero76 cells (hCoV-19/Germany/Muenster_FI1103201/2020, GISAID accession EPI_ISL_463008) was isolated at the Institute of Virology, Muenster, Germany, from a patient returning from the Southern Tyrolean ski areas. Infectious virus stocks were propagated in Vero-TMPRSS2 cells. The SARS-CoV-2 strain used to infect hamsters (SARS-CoV-2/human/BEL/GHB-03021/2020, GISAID accession EPI_ISL_407976) was isolated at the Rega Institute, Leuven, Belgium from a nasopharyngeal swab taken from an RT-qPCR-confirmed asymptomatic patient returning from Wuhan, China in February, 2020(Spiteri *et al*). Infectious virus stocks were propagated in Vero E6 cells.

### Hamster model and inoculation

The hamster model of acute SARS-CoV-2 infection has been previously described(Kaptein *et al*, 2020). The hamsters were purchased from Janvier Laboratories. They were housed in ventilated isolator cages (IsoCage N Biocontainment System, Tecniplast) at two per cage with ad libitum access to food and water and cage enrichment (a wood block). Housing conditions and experimental procedures were approved by the ethics committee for animal experimentation of KU Leuven (License P065-2020). In brief, wild-type female golden Syrian hamsters (*Mesocricetus auratus*) six to ten weeks of age were inoculated intranasally with 50 μL solution containing 2×10^6^ TCID_50_ of SARS-CoV-2. Four days post-inoculation the hamsters were euthanized and whole lungs were collected and subjected to RNA isolation as described under ‘RNA extraction and DNase treatment’.

### Cell culture and inoculation

Vero76 cells (African green monkey kidney, ATCC CRL-1587) were cultured in Dulbecco’s Modified Eagle Medium (Gibco, 41965-039) supplemented with 10% Fetal Bovine Serum (Gibco, 10270-106) and 1% penicillin/streptomycin (Gibco, 10378016) in a humidified atmosphere at 37°C under 5% CO_2_. Cells were routinely checked for mycoplasma contamination with the MycoAlert mycoplasma detection kit (Lonza, LT07-318). Cells were authenticated by STR-typing, amplification, and sequencing. Vero76 cells were inoculated with SARS-CoV-2 at MOI 0.001. For this, they were incubated for 1 h at 37°C under 5% CO_2_ with an equal volume of diluted infectious stock. Then the virus-containing medium was removed, the cells were washed twice with PBS, and standard culture medium was added. Twenty-four hours post-inoculation, both the culture supernatant and the cells were harvested and subjected to RNA isolation as described under ‘RNA extraction and DNase treatment’.

### Stable short hairpin RNA knockdown

Stable FTO-depleted Vero76 cell lines were generated by lentiviral infection. Briefly, the FTO-targeting and Control shRNAs were designed *in silico* with Blastn (NCBI) and siRNA wizard (Invivogen) online tools. They are as follows: FTO (shFTO-sense: 5ʹ-GAA CAA GCC CAG CAT GTG A-3ʹ, shFTO-antisense: 5ʹ-T CAC ATG CTG GGC TTG TTC-3ʹ), Control (shCTR-sense: 5ʹ-GAC GCA ACC TGA GGA CTA A-3ʹ, shCTR-antisense: 5ʹ-T TAG TCC TCA GGT TGC GTC-3ʹ). They were cloned into the pSUPER.retro.puro vector (OligoEngine, VEC-PRT-0002), and packaged into lentiviruses following pVPack-VSV-G vector (Agilent, 217567) co-transfection in 293GP cells with TransIT-LT1 Transfection Reagent (Mirus, 2300). Two days later, the resulting filtered lentivirus-containing medium was used to infect Vero76 cells, which were further selected with puromycin at 10 μg/mL. Knockdown efficiency was validated at both the RNA and protein levels.

### Western blotting

To assess the FTO protein level after shRNA knockdown, cells were harvested and lysed in lysis buffer (50 mM Tris-HCl pH 7.5, 150 mM NaCl, 1 mM EDTA, 0.5% NP-40) supplemented with proteinase and phosphatase inhibitors. Cell debris were removed from the lysate by centrifuging at 14,000 g and 4°C for 10 min. Protein concentration was measured with DC protein assay reagents (BioRad, 5000116). protein lysate (30 μg) was loaded onto and run through a 10% SDS-PAGE gel and transferred to a PVDF membrane (Merck, IPVH00010). Membranes were blocked with 5% (w/v) nonfat dry milk (Cell Signaling, 9999S) in PBST for 1 h at room temperature and then immunoblotted overnight at 4 °C with corresponding primary antibodies: anti-FTO (Abcam, ab124892, 1/1000) or anti-β-Actin (Sigma, A5316, 1/1000). Membranes were washed three times with PBST for 5 min and incubated for 1 h at room temperature with HRP-conjugated secondary anti-rabbit IgG (GE Healthcare, NA934, 1/2000) or anti-mouse IgG (Amersham, NXA931V, 1/2000). Membranes were finally washed three times with PBST for 5 min and protein levels were visualized by enhanced chemiluminescence (Perkin Elmer, NEL104001EA) according to the manufacturer’s instructions. Membranes were finally imaged and quantified with Image Lab software (Version 6.1). β-Actin was used as a loading control.

### RNA extraction and DNase treatment

Total RNA from infected cells, their supernatants, and hamster lung tissues were extracted with Trizol (Invitrogen, 15596026) according to the manufacturer’s instructions. Contaminating DNA was removed from supernatants with the DNA-free DNA Removal Kit (Invitrogen, AM1906) and from infected cells and hamster tissues with the RNase-Free DNase set coupled with the RNeasy Midi Kit (Qiagen, 79256 & 75144), according to the manufacturers’ protocols. RNA concentration was assessed with a Qubit 4 Fluorometer (Invitrogen, Q33226).

### RT-qPCR

To assess levels of m^6^A machinery component mRNAs upon SARS-CoV-2 infection, isolated total RNA from both non-infected and infected Vero76 cells was converted to cDNA with the SuperScrip II Reverse Transcriptase (Invitrogen, 18064-014) with a random primer (Invitrogen, 48190011). All PCR reactions were carried out with the LightCycler 480 SYBR Green I Master (Roche, 04887352001) according to the manufacturer’s protocol and amplification was quantified on an LightCycler 480 II Real-Time PCR System (Roche). In all cases, average threshold cycles were determined from at least triplicate reactions and relative gene expression levels were calculated by the 2-ΔΔCt method(Livak & Schmittgen, 2001). RNA expression levels were normalized to the mean expression of housekeeping genes (GAPDH and Tubulin). The primer pairs used for qPCR are listed in **Supplementary Table 1**.

### m^6^A sequencing (m^6^A-seq)

m^6^A-seq was performed as previously described with some modifications(Dominissini *et al*, 2012). Briefly, total RNAs (viral RNA/cellular RNA ratio = 1/10) were fragmented by mixing 90 µL RNA (at 0.9 µg/µL) with 10 µL of 10x fragmentation buffer (100 mM Tris-HCl and 100 mM ZnCl2). The mix was incubated at 94°C for 40 sec in a preheated thermocycler with the lid open. The reaction was stopped with 10 µL of 0.5 M EDTA, the mixture was incubated on ice, and RNA fragments were precipitated overnight at −80°C. RNA fragment sizes were assessed on an RNA 6000 Nano chip (Agilent, 5067-1511) with a 2100 Bioanalyzer (Agilent). 5 µg fragmented RNA was kept at −80°C as input. m6A immunoprecipitation was performed as follows: 250 μg fragmented RNA was denatured by heating at 70°C for 5 min and placed on ice. Denatured RNA was then mixed with 7 μg anti-m^6^A antibody (Synaptic System, 202003) in 500 µL of 1x IP buffer (10 mM Tris-HCl, 150 mM NaCl, 0.1% IGEPAL CA-630) supplemented with 10 µL cOmplete EDTA-free Protease Inhibitor Cocktail (Roche, 11873580001), 5 µL of 200 mM Ribonucleoside Vanadyl Complex (RVC) (Sigma, R3380), and 200 U RNasin Ribonuclease Inhibitors (Promega, N2511). The mix was incubated overnight at 4°C with rotation. 50 µL Dynabeads Protein G (Invitrogen, 10004D) were washed twice with 1x IP buffer supplemented with protease inhibitor and blocked in IP buffer supplemented with 0.5 mg/mL BSA (Sigma, A7284) for 1 h on a rotation wheel. Beads were washed twice, added to the immunoprecipitation mix, and incubated for 2 h at 4°C with rotation. After three washes with IP buffer containing RNasin and RVC, the beads were resuspended in 200 µL 1x IP buffer and m^6^A-containing RNA fragments were eluted with TriPure Isolation Reagent (Roche, TRIPURE-RO). After RNA recovery, cDNA libraries were prepared with the SMARTer Stranded Total RNA-Seq Kit v2 - Pico Input Mammalian (Takara, 634413). All samples were sequenced on Illumina NextSeq500 (Illumina).

### m^6^A RIP-qPCR

To assess m^6^A enrichment on SARS-CoV-2 in the hamster model, m^6^A-RIP-qPCR was performed as previously described above for m^6^A-seq, with some modifications. First, qPCR primers amplifying peak regions identified on SARS-CoV-2 in the Vero76 model were designed and synthesized. Additionally, a positive control consisting of a 1089-nucleotide methylated transcript (Renilla luciferase) was *in vitro* transcribed with the MEGAscript T7 Transcription Kit (Invitrogen, AM1334) according to the manufacturer’s instructions, with addition of m^6^ATP (TriLink, N-1013) instead of ATP nucleotides in the reaction mix. The resulting m^6^A transcript was then mixed at ratio 1/500 with 200 μg total RNA from infected hamster lung tissues and m^6^A-IPed as described above. An equal amount of anti-IgG antibody (Merck, PP64) was also used for immunoprecipitation. Input RNA, IgG- and m^6^A-IPed RNA were subjected to reverse transcription and subsequent qPCR. Enrichment of each IP in each peak region was calculated by normalizing to input. The viral ORF1a, ORF1b, and spike regions were used as negative controls. The methylated control and the primer sequences used are provided in **Supplementary Table 1**.

### Read pre-processing

Sequencing data were pre-processed in the following steps. First, the raw sequencing data were analyzed with FastQC (v0.11.5). Low-complexity reads were removed with the AfterQC tool (v0.9.6) with default parameters. To get rid of reads originating from rRNA or tRNA, reads were mapped with Bowtie2 (v2.3.4.1) to Chlorocebus sabaeus tRNA and rRNA sequences downloaded from Ensembl (v101). Unmapped reads were then further processed with Trimmomatic (v0.33) using default parameters to remove adapter sequences. The resulting fastq data were again analyzed with FastQC to ensure that no further processing was needed. Transcript sequences from Vero76 cells were downloaded from NCBI using with search parameters for Chlorocebus sabaeus. A metagenome built from the Chlorocebus sabaeus (Ensembl v100-ChlSab1.1.dna_sm.toplevel.fa.gz) and SARS-CoV-2 (NC_045512.2) reference genomes was created as extra chromosome. The pre-processed reads were further mapped to the constructed metagenome with the STAR algorithm (v2.6.0c) using the Ensembl reference transcriptome (v100) for Chlorocebus sabaeus and SARS-CoV-2 transcriptome from Miladi and al (Miladi *et al*, 2020).

### m^6^A-seq analysis

Reads localized to the SARS-CoV-2 extra chromosome were extracted with grep (v2.25) from bam files. m^6^A peak regions were identified by applying the m6aViewer peak-calling tool (v1.6.1) on immunoprecipitated (IP) samples, using their input counterpart to estimate background noise (corrected p-value < 0.05). Peaks were resized to 100 bp on both sides of the identified summit. To obtain visual representations of local enrichment profiles, normalized (HPB) bedgraph files were generated for each input and IP sample using bamTobw (https://github.com/YangLab/bamTobw) and uploaded into the IGV tool (v2.9.4). Enrichment ratios were defined for each condition as IP over input peak height ratio. Peaks further considered in the analysis were only the ones significant in all the replicates.

In order to validate our m^6^A detection method, we also analyzed the peaks detected on the Chlorocebus sabaeus part of the metagenome. Known conserved peaks in human and mouse were compared with those found in Chlorocebus sabaeus.

To perform the motif analysis, FIMO from the meme-suite (http://meme-suite.org) was used to screen for m^6^A consensus motifs such as RRACH, UGAC, and MGACK within the identified peak regions (p-value < 0.01). Motif densities were performed with the density function of R software (v4.1.2).

### Viral titration assay

SARS-CoV-2 infectious titers were measured by standard viral plaque assay as previously described(Hoffmann *et al*, 2021). 50,000 control or FTO-depleted Vero76 cells were seeded into a 48-well plate. Twenty-four hours later, the cells were infected with SARS-CoV-2 at MOI 0.001 and incubated for 1 h at 37°C, 5% CO_2_. Then the inoculum was discarded, the cells were washed twice with PBS, and placed in DMEM supplemented with 2% instead of 10% FBS. Infected cells were cultured for three days post-inoculation. Each day, culture supernatants were harvested and confluent Vero-TMPRSS2 cells were infected with 10-fold serial dilutions of supernatant for 1 h at 37°C. Thereafter, the infection medium was discarded and the cells were incubated with culture medium containing 1% (w/v) methyl cellulose. Plaques were counted 45 h post-inoculation and the infectious titer was determined in log_10_ plaque-forming units per mL (log_10_ (pfu/mL)).

### Phylogenetic analysis on human coronaviruses’ Nucleocapsid

Nucleotide sequences of the nucleocapsid and TRS_L regions of SARS-CoV-2 (NC_045512.2:28274-29533), SARS-CoV (NC_004718.3:28120-29388), MERS-CoV (NC_019843.3:28566-29807), HCoV-HKU1 (NC_006577.2:28320-29645), HCoV-OC43 (NC_006213.1:29079-30425), HCoV-229E (NC_002645.1:25686-26855), HCoV-NL63 (NC_005831.2:26133-27266) were obtained from the National Center for Biotechnology Information (NCBI) Reference Sequence (RefSeq) database. Multiple sequence alignments were performed with Geneious Prime (v2021.1.1) with Muscle default parameters (v*3.8.425*) (Song *et al*, 2020; Gong *et al*, 2020). Maximum likelihood phylogenetic trees were generated with Geneious Prime (v.10.2.4), with the Unweighted Pair Group Method with the Arithmatic Mean (UPGMA) model and 1,000 bootstrap replicates. Nucleotide similarities were obtained using SimPlot (v.3.5.1) with parameters set at window 200 bp or 60 bp, step 20 bp, GapStrip: on, Kimura model.

### Mutation analysis of SARS-CoV-2 isolate nucleocapsids

Variation annotations for SARS-CoV-2 nucleocapsid and TRS_L sequences were downloaded from the 2019 Novel Coronavirus Resource (https://ngdc.cncb.ac.cn/ncov/) provided by the National Genomics Data Center, China National Center for Bioinformation at the Beijing Institute of Genomics, Chinese Academy of Sciences on the 25^th^ April 2022. This data set represents variations from a total of 9,557,377 isolated SARS-CoV-2 variants with complete sequences from different countries located in six geographic areas. This included 5,161,011 sequences from Europe, 3,317,560 from North America, 733,146 from Asia, 197,107 from South America, 78,912 from Africa, and 69,641 from Oceania. Additionally, lineage information on these different sequences was also retrieved and lineage distribution assessed. Mutation frequencies were calculated from the number of isolates with variation and the total number of SARS-CoV-2 isolates worldwide.

### Patient data collection

Single-cell RNA-seq data were retrieved from Wauters et al.(Wauters *et al*, 2021). The patient cohort was composed of 13 non-COVID-19 and 22 COVID-19 pneumonia patients, collected from the University Hospitals Leuven, between March 31st 2020 and May 4th 2020, and processed scRNA-seq data on COVID-19 BAL fluid from 9 patients by Liao et al. (Liao *et al*, 2020). Disease severity was defined as ‘mild’ or ‘severe’ on the basis of the level of respiratory support at the time of sampling. Specifically, patients with ‘mild’ disease required no respiratory support or supplemental oxygen through a nasal cannula, whereas patients with ‘severe’ disease were mechanically ventilated or received extracorporeal membrane oxygenation. For analysis, we retrieved normalized gene counts of our genes of interest from an object analyzed with the Seurat package (v3.1.4). Cells of either control or COVID-19 samples were additionally defined according to their SARS-CoV-2 expression status (at least one read on SARS-CoV-2 genome required for positive status). For the classifier analysis we filtered the dataset in order to keep the epithelial cells and more precisely the AT2, ciliated, secretory, and Hillock-type cells.

RNA-seq data from nasal swab samples were retrieved from Ng et al. (GSE163151) (Ng *et al*, 2021). The patient cohort includes 93 control patients, defined by non-pathogenic status, and 147 SARS-CoV-2-positive patients. Only SARS-CoV-2-positive patients associated with severity status were considered, amounting to 80 COVID-19 patients. Specifically, patients with ‘severe’ disease required intensive care unit (ICU) support, whereas patients with ‘mild’ disease were either outpatients or non-ICU hospitalized. For analysis, gene expression levels in transcripts per million (TPM) were calculated from the raw counts using gene sizes from ensembl human transcriptome (v92).

RNA-seq data from blood samples were retrieved from Overmyer et al. (GSE157103) (Overmyer *et al*, 2021). The patient cohort consists of 26 non-COVID-19 and 100 COVID-19 patients. Disease severity was defined as ‘mild’ or ‘severe’ on the basis of ICU need. Specifically, ‘severe’ disease required ICU support, whereas ‘mild’ patients were non-ICU hospitalized. TPM gene expression levels were directly used to build the classifier.

### Severity classifier construction

Classifiers of COVID-19 severity status were built with a K-Nearest Neighbors (k-NN) model approach. In brief, to distinguish mild from severe COVID-19 patients, a k-NN algorithm (as implemented in the R package FNN (v1.1.3)) was trained with gene-of-interest expression levels from the above-mentioned datasets. A hundred bootstrap replicates of tri-fold cross-validation were performed to reduce model error and obtain a better estimate of the Area Under the Receiver Operating Characteristic (ROC) Curve (AUC). Results were represented as ROC curves by means of the R packages pROC (v1.18.0). FTO performance as a severity classifier was benchmarked against expression of well-known cytokines and interferon-stimulated genes. Among these, we used the interferon-stimulated gene IFI6, which is strongly induced by SARS-CoV-2 in upper airway cells, the pro-inflammatory cytokines IL1B, IL1R2 and IL6, whose production is increased in macrophages during severe COVID-19, and the chemokine receptors CCR2 and CCR5, mediating monocyte infiltration in inflammatory diseases(Mick *et al*, 2020; Merad & Martin, 2020).

### Statistical analysis

All statistical analyses were performed with either GraphPad Prism 9 or the computing environment R (v4.1.2). Unless otherwise specified, all statistics were evaluated by a two-tailed unpaired Student’s t-test with *p* < 0.05 as statistical significance criterion. Data are reported as means ± SEM of at least two biological independent experiments conducted with technical replicates. One-tailed Pearson’s test was used for the correlation analyses.

### Data availability

The m^6^A-seq data supporting the findings of this study have been deposited in the GEO repository under accession code “GSE201626“. Datasets supporting our study (m^6^A-seq, coronaviruses sequences, SARS-CoV-2 mutations, BAL scRNA-seq and bulk RNA-seq) are either publicly available or published data and sources are provided in this paper.

## Acknowledgments

Figures were created using Biorender.com. We thank Tina Van Buyten for her excellent technical assistance with the homogenization of the hamster lung samples. L.M. and M.C. were supported by a Télévie grant. M.B., B.H., P.P., E.C., R.D. and J.J. were supported by the Belgian FNRS. F.F.’s lab was funded by grants from the FNRS, the ‘Action de Recherche Concertée’ (ARC; AUWB-2018-2023 ULB-No 7), an FNRS Welbio grant, the ULB Foundation. This work was supported by the Belgian FNRS ‘Crédit Urgent de Recherche’ (CUR, Ref. 40002805). The computational resources and services used in this work were provided by the VSC (Flemish Supercomputer Center), funded by the Research Foundation - Flanders (FWO) and the Flemish Government – department EWI.

## Author contributions

L.M., L.N., R.D., L.D., J.J. and F.F. initiated the work. L.M., M.C., M.B., J.J. and F.F. designed the experiments and interpreted the data. L.M., B.H. and P.P. performed the FTO RNAi experiments in Vero cells. H.H.W. and S.P. helped with Vero cells infection and viral titer assays. R.A. and L.D. provided infected hamsters’ lung tissues. E.C. performed m^6^A RIP-qPCR and m^6^A deep-sequencing experiments. B.B. and D.L. provided data and technical support on scRNA-seq analysis. L.M., M.C. and M.B. performed the bioinformatics analyses. J.J. and F.F. directed the study. L.M., M.C. and M.B prepared the figures. L.M., M.C, M.B, J.J., and F.F. wrote the manuscript.

## Supplementary Materials

**Fig. S1.**
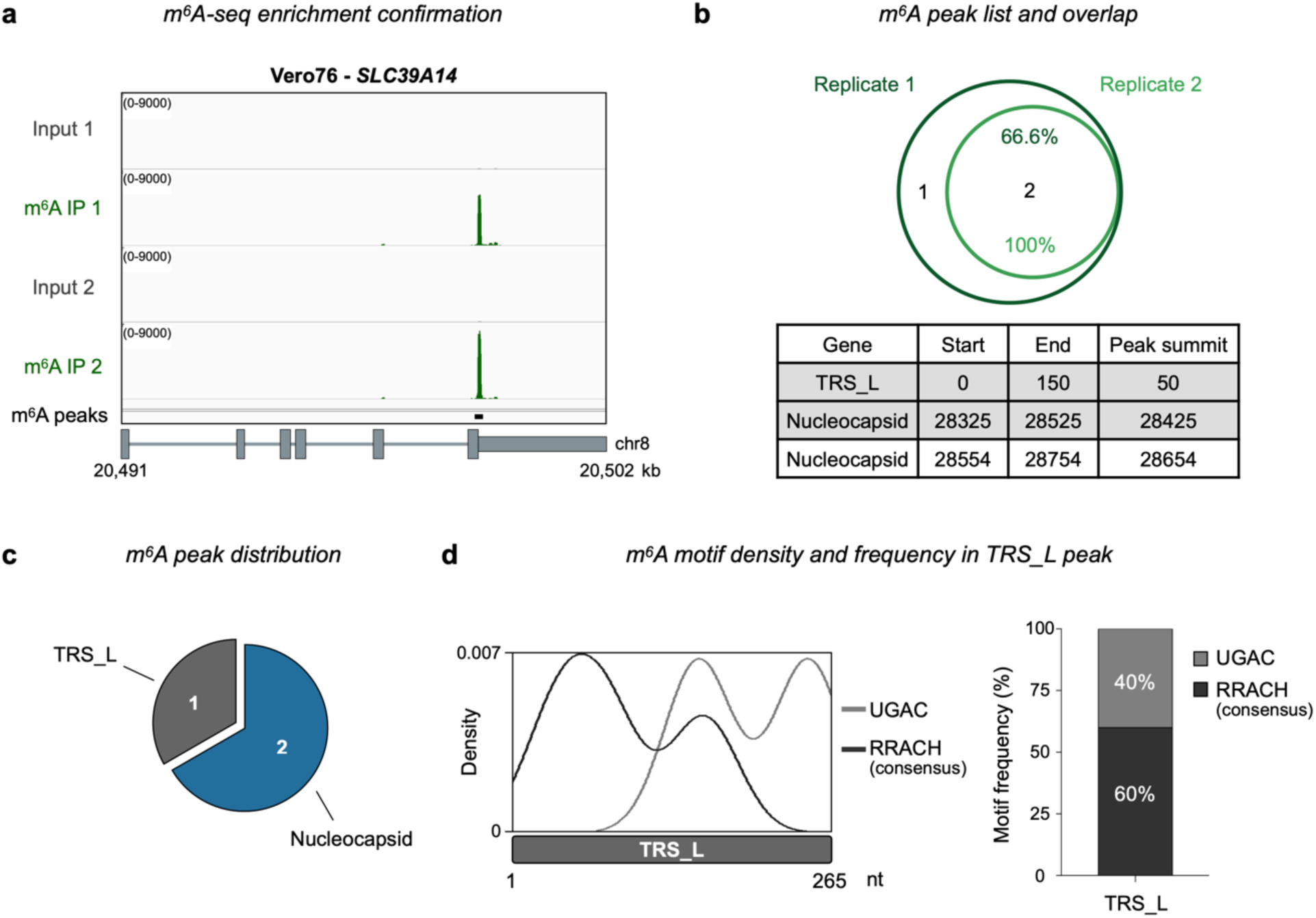
m^6^A decorates SARS-CoV-2 RNA *in vitro* and *in vivo*. **a** IGV tracks display an exemplative m^6^A-seq profile of the orthologous gene *SLC39A14* as a positive control for enrichment through m^6^A-IP. Input and m^6^A IP from two biological replicates are labeled in gray and green, respectively. Black squares represent m^6^A peaks. Gene architecture is shown beneath. **b** Venn diagram (upper panel) showing the between-replicate overlap of m^6^A peaks identified by m^6^A-seq in Vero76 cells. Peaks are listed below the diagram. Rows colored in gray highlight m^6^A peaks shared by both biological replicates. **c** Pie chart illustrating the number and distribution of all m^6^A peaks identified along SARS-CoV-2 RNA. **d** Density plot (left panel) showing known m^6^A motifs (RRACH, UGAC) across the m^6^A peak in the SARS-CoV-2 TRS_L region. Stacked bar chart (right panel) showing motif frequencies within the m^6^A peak in the SARS-CoV-2 TRS_L region.

**Fig. S2.**
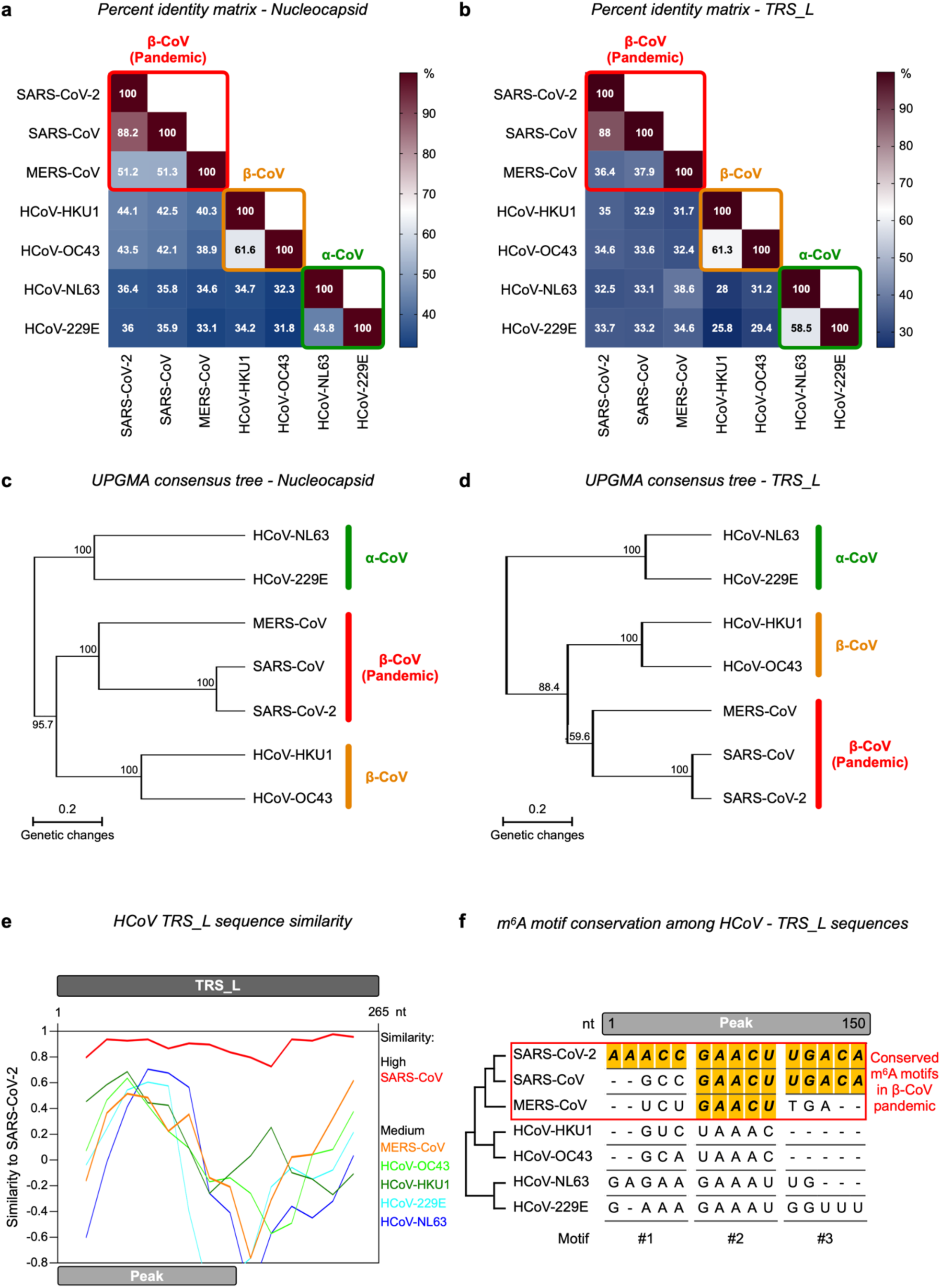
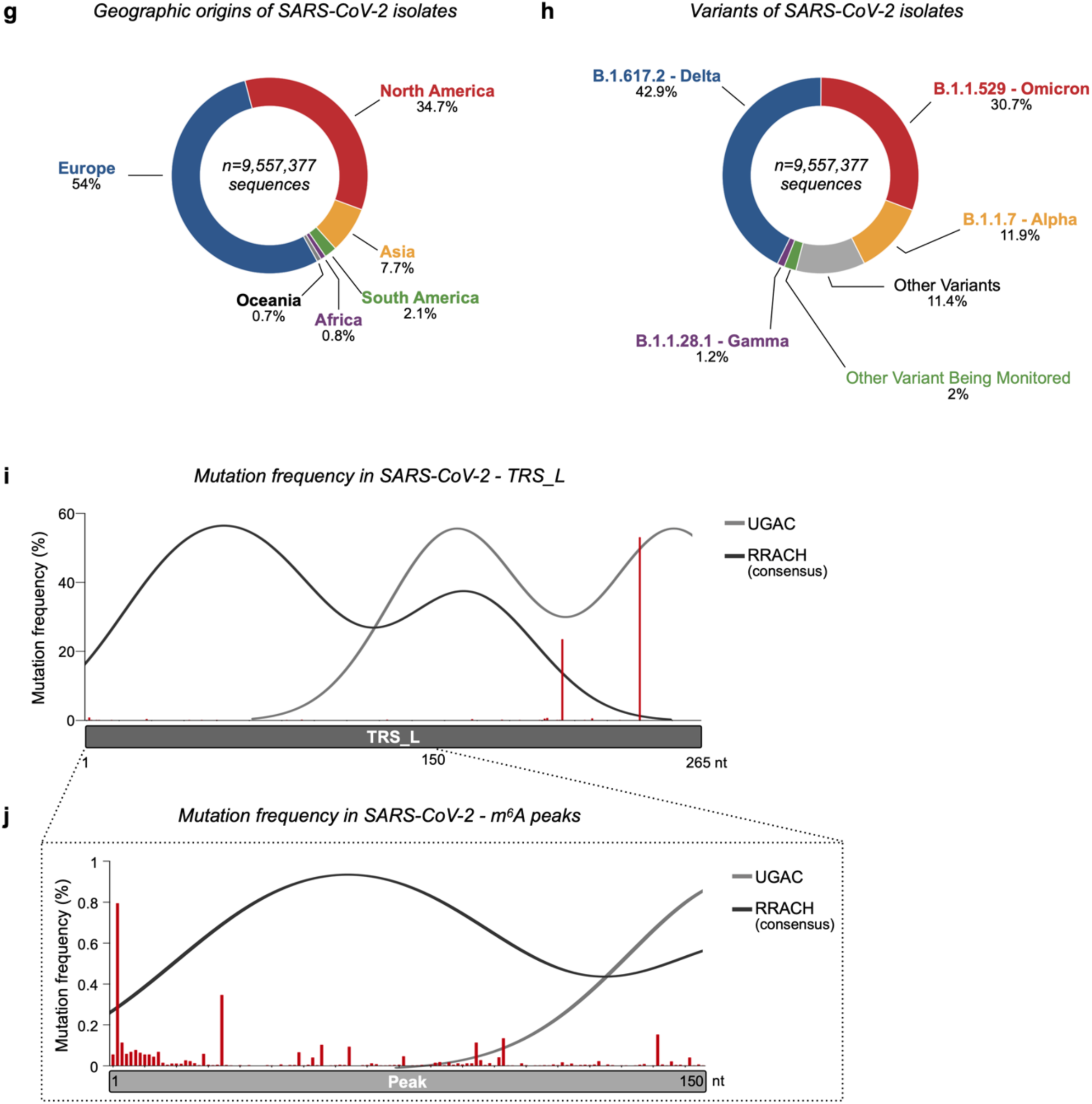
m^6^A sites are specific and conserved among SARS-CoV-2 strains. **a**, **b** Percent identity matrix of nucleocapsid (a) and TRS_L (b) sequences of human coronaviruses (HCoV), created with Geneious Prime, using the Muscle Multiple Sequence Alignment (MSA) tool and default parameters. Three groups are indicated: pandemic β-CoV (SARS-CoV-2, SARS-CoV, and MERS-CoV), β-CoV (HCoV-HKU1, HCoV-OC43), and α-CoV (HCoV-NL63, HCoV-229E). **c**, **d** Maximum likelihood phylogenetic tree of nucleocapsid (c) and TRS_L (d) sequences of human coronaviruses, generated using Geneious Prime with the Unweighted Pair Group Method with Arithmatic Mean (UPGMA) model and 1,000 bootstrap replicates. Numbers on the branches indicate bootstrap support values. Branch scale bars are shown as 0.2 substitutions per site. **e** TRS_L-region sequence similarity among human coronaviruses. This plot is based on the full-length TRS_L sequence, derived through MSA of all human coronaviruses with the following parameter settings: window 60 bp, step 20 bp, GapStrip: on, Kimura model. **f** MSA describing m^6^A consensus motif variations across human coronavirus TRS_L sequences within the peaks identified in SARS-CoV-2. m^6^A consensus sequences conserved across HCoVs are colored in yellow. **g** Pie chart showing the number of SARS-CoV-2 sequences available from the NGDC-CNCB as of April 25^th^ 2022 for six different regions: Europe (blue, 54%), North America (red, 34.7%), Asia (yellow, 7.7%), South America (green, 2.1%), Africa (purple, 0.8%) and Oceania (gray, 0.7%). **h** Pie chart displaying the number of SARS-CoV-2 sequences from different lineages available from the NGDC-CNCB as of April 25^th^ 2022. Only variants with a frequency higher than 1% are displayed. Variants with lower frequency are either grouped into the “Other Variants Being Monitored” category (Iota, Beta, Epsilon, Mu, Eta, Kappa, Zeta, and Theta) or the “Other Variants” category. **i** Bar chart displaying, for a total of 9,557,377 SARS-CoV-2 complete isolates, the mutation frequency in TRS_L regions across SARS-CoV-2 variants as of April 25^th^ 2022. **j** Bar chart (red) and density plot (gray) displaying, across SARS-CoV-2 variants, the mutation frequency and distribution of known m^6^A motifs (RRACH and UGAC) across m^6^A peaks identified within the TRS_L.

**Fig. S3.**
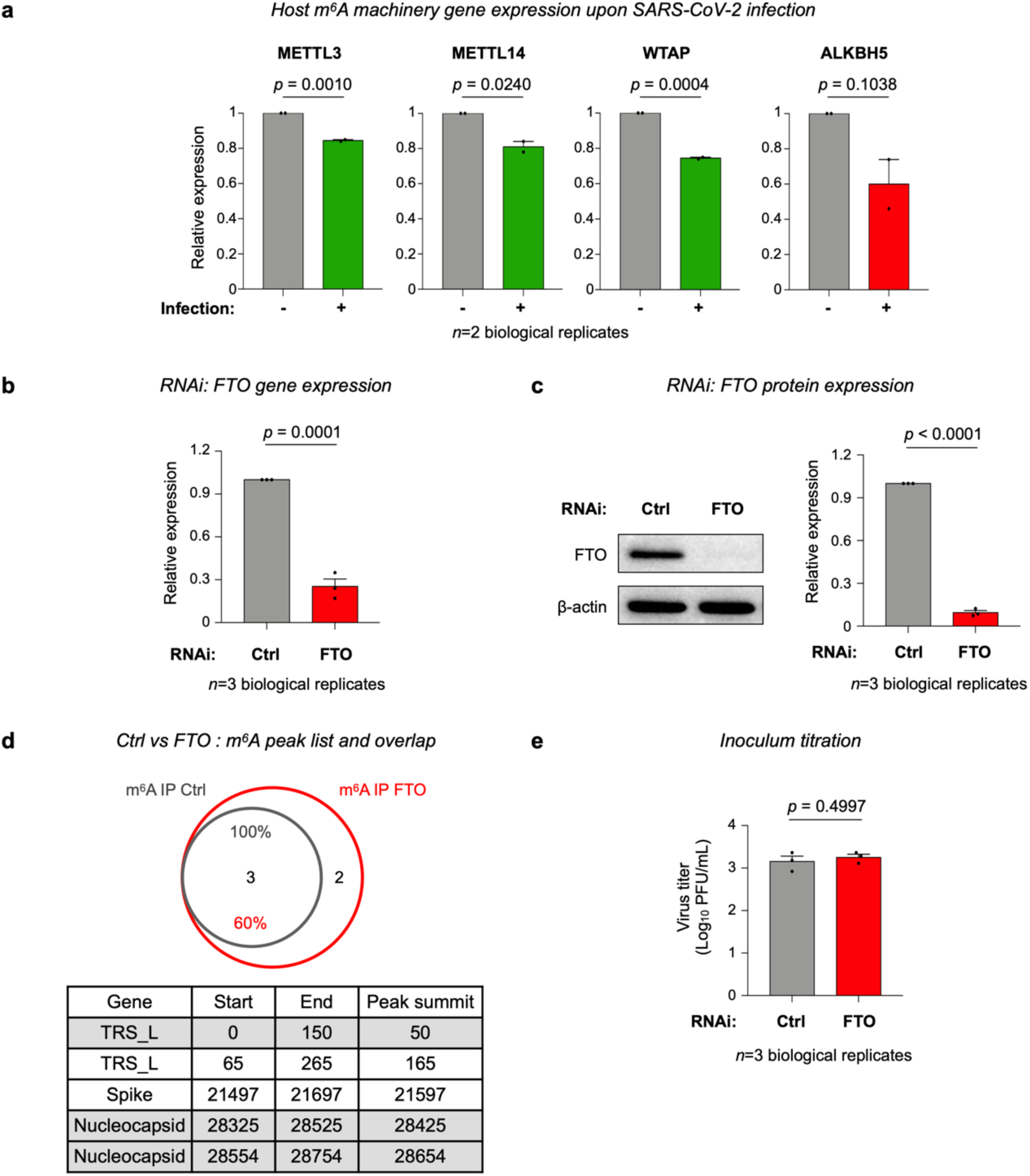
The m^6^A demethylase FTO impairs SARS-CoV-2 replication. **a** Host m^6^A machinery gene expression upon SARS-CoV-2 infection in monkey cells. RT-qPCR assays displaying altered levels of mRNAs encoding host m^6^A machinery components after SARS-CoV-2 infection of Vero76 cells. Dots represent mean expression for independent experiments conducted with technical triplicates. **b** RT-qPCR assay validating decreased mRNA-level *FTO* expression upon FTO depletion by RNAi in Vero76 cells. Dots represent mean expression for three independent experiments conducted with technical triplicates. **c** Western blot assays confirm a decreased FTO protein level upon FTO depletion by RNAi in Vero76 cells. Left panel: representative western blot, β-actin serving as a protein loading control. Right panel: western blot quantification from three biological replicates. **d** The Venn diagram (upper panel) shows the overlap between m^6^A peaks identified by m^6^A-seq in infected controls (gray) and FTO-depleted Vero76 cells (red). All m^6^A peaks identified in both control and FTO-depleted Vero76 cells are listed below the diagram. Rows colored in gray highlight m^6^A peaks shared by both controls and FTO-depleted cells. e Bar chart showing the SARS-CoV-2 inoculum used to infect control (gray) and FTO-depleted (red) Vero76 cells. Plaque-forming units (PFU) of infectious SARS-CoV-2 particles in culture supernatants were determined by plaque titration with Vero-TMPRSS2 cells. Dots represent mean expression for three independent experiments conducted with technical triplicates. Data represent means ± SEM. *P*-values were determined with a two-tailed unpaired Student’s t test.

**Fig. S4.**
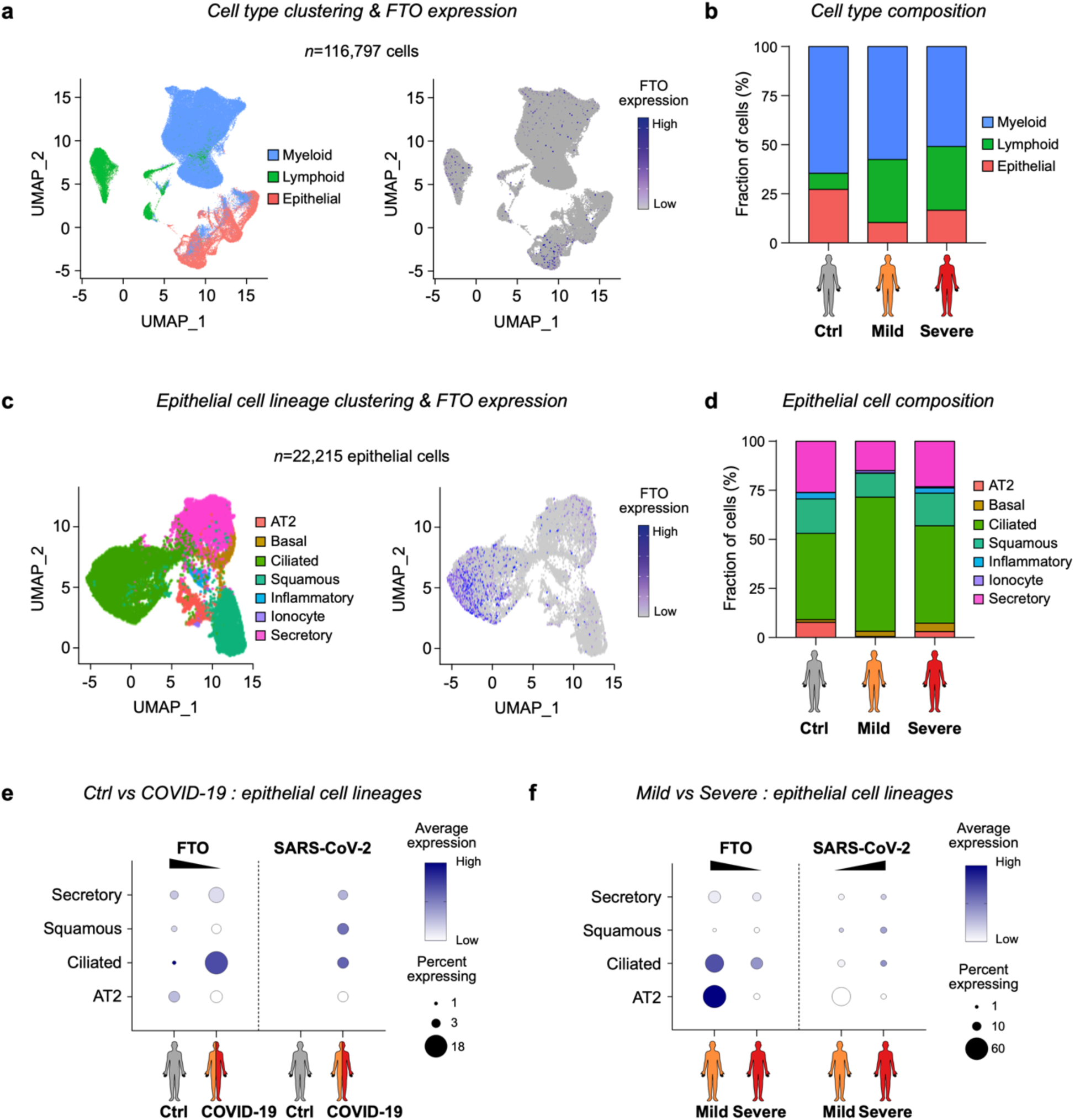
*FTO* downregulation correlates with higher SARS-CoV-2 expression. **a** UMAP representation (left panel) showing scRNA-seq-based clustering of the 116,797 cells present in BAL samples from all patients. Each dot represents one cell and is color-coded for the indicated cell domain: myeloid (blue) lymphoid (green), or epithelial (pink). UMAP representation (right panel) displaying *FTO* expression in the different cell domains identified in BAL datasets. **b** Bar plot showing the differential cell-type distribution of BAL datasets from controls and COVID-19 patients with mild and severe symptoms. **c** UMAP representation (left panel) illustrating scRNA-seq-based clustering of the 22,215 epithelial cells in BAL samples from all patients. Each dot represents one cell and is color-coded for the indicated cell lineage: AT2 (salmon), basal (yellow), ciliated (green), squamous (teal), inflammatory (blue), ionocyte (purple), or secretory (pink). UMAP representation (right panel) showing *FTO* expression in the different cell lineages identified in BAL datasets. **d** Bar plot displaying differential cell-lineage composition in BAL datasets from control patients and COVID-19 patients with mild or severe symptoms. **e, f** Dot plots showing differential expression of the *FTO* gene and of SARS-CoV-2, in different epithelial-cell lineages identified in BAL datasets, between control and COVID-19 patients (e) and between patients with mild and severe COVID-19 (f). In the gene-expression dot plots, dot size represents the proportion of cells of a particular cell type expressing the gene of interest and dot color represents the average gene expression level in that cell type.

**Fig. S5.**
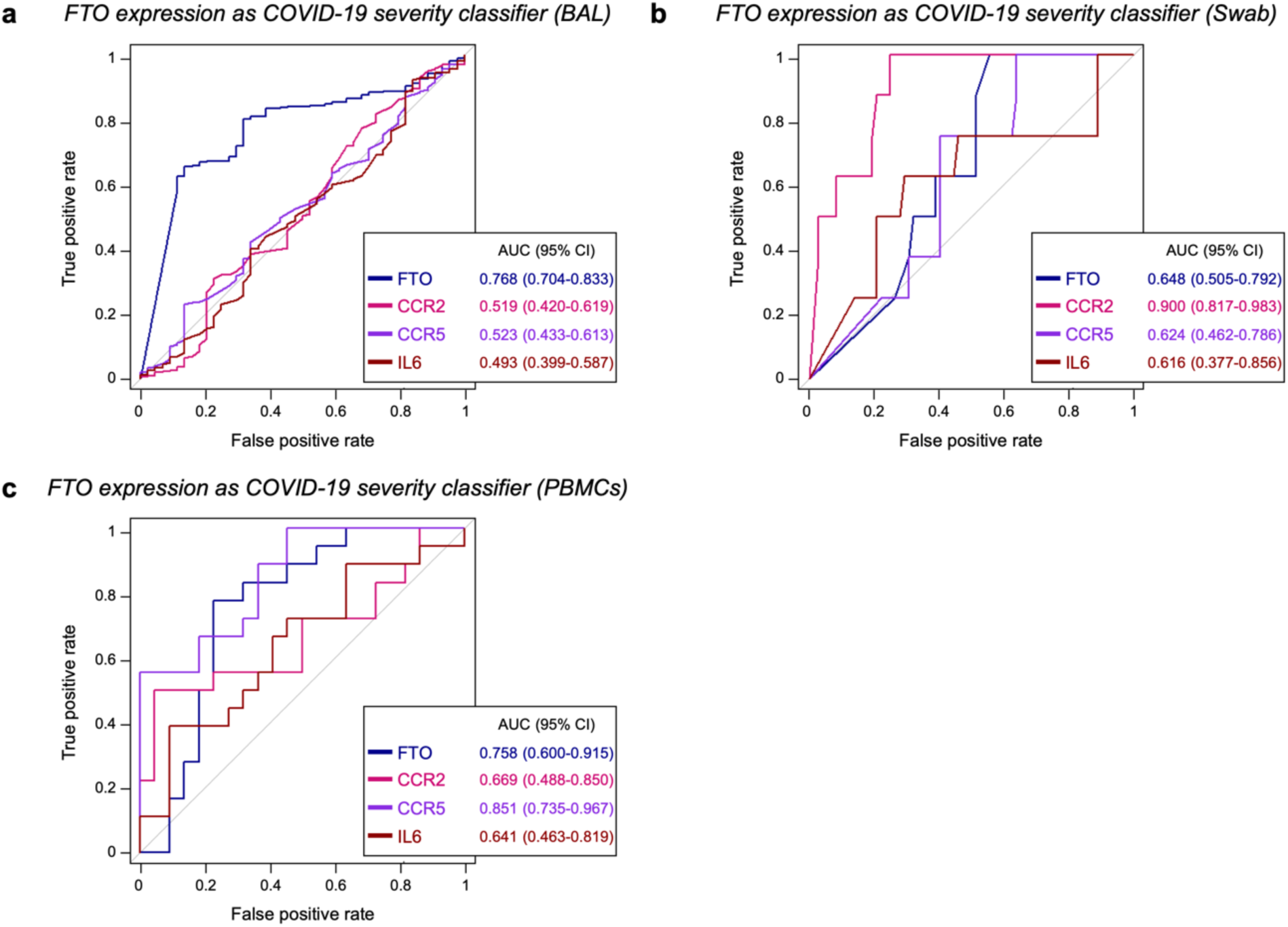
*FTO* expression in patient samples classifies COVID-19 severity. **a, b**, **c** Receiver Operating Characteristic (ROC) curves showing the performance, as a diagnostic COVID-19 severity classifier, of FTO expression versus CCR2, CCR5, and IL6 expression in patient epithelial cells from BAL samples (a), patient cells from nasal swab samples (b), and patient PBMCs from blood samples (c). The mean area under the ROC curve (AUC) and the 95% confidence interval are indicated.

**Table. S1.**
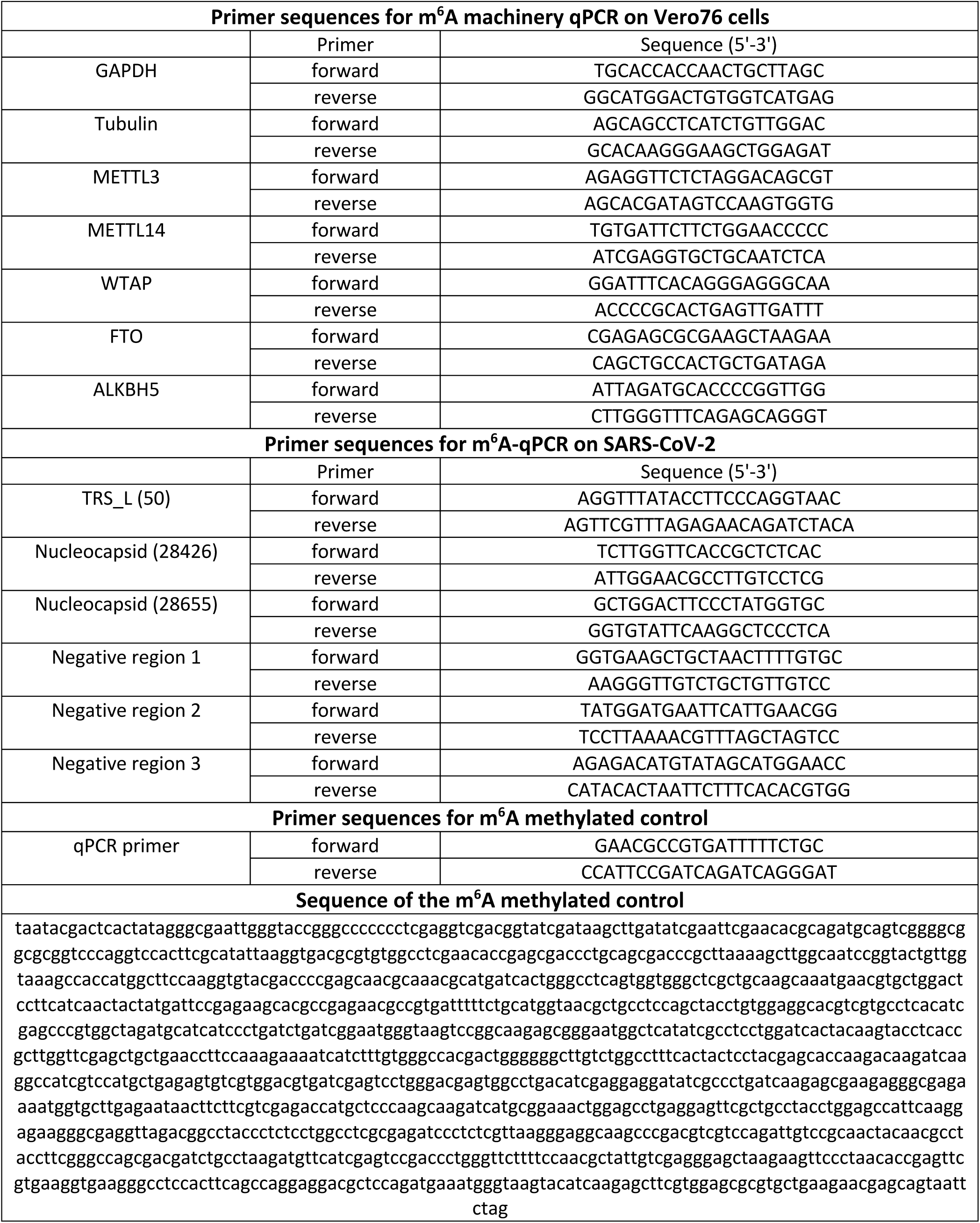
Sequences information of oligo and primers used in this study.

